# Transcriptome and fatty-acid signatures of adipocyte hypertrophy and its non-invasive MR-based characterization in human adipose tissue

**DOI:** 10.1101/2021.11.20.468818

**Authors:** Julius Honecker, Stefan Ruschke, Claudine Seeliger, Samantha Laber, Sophie Strobel, Priska Pröll, Christoffer Nellaker, Cecilia M. Lindgren, Ulrich Kulozik, Josef Ecker, Dimitrios C. Karampinos, Melina Claussnitzer, Hans Hauner

## Abstract

Adipose tissue is an organ with great plasticity and its hypertrophic expansion is associated with adipocyte dysfunction. How changes in adipocyte morphology are linked to gene expression and which cellular functions are affected remains elusive. We show that adipocyte hypertrophy is associated with transcriptomic changes using RNA-Seq data obtained from adipose tissue and size-separated adipocytes. Genes involved in oxidative phosphorylation, protein biosynthesis and fatty acid metabolism were down-regulated in large adipocytes while genes involved in inflammation were upregulated. For mitochondrial function, a reduction in the expression of thermogenesis related genes and estimated brown/beige adipocyte content was observed in individuals with large adipocytes. As a novel finding the total adipose tissue fatty acid composition was dependent on cell size and depot. MR spectroscopy methods for clinical scanning were developed to characterize adipocyte size and fatty acid composition in a fast and non-invasive manner. Together, the present data provides mechanistic insights on how adipocyte hypertrophy contributes to the manifestation of metabolic disease at the molecular and cellular level. MR spectroscopy was identified as a promising technique for an in-parallel assessment of adipose morphology and fatty acid composition allowing to translate our findings into an improved, non-invasive phenotyping of adipose tissue.

## INTRODUCTION

A higher body fat percentage and thus an expanded white adipose tissue (WAT) mass is a central hallmark of obesity. An increase in the number of small adipocytes (hyperplasia) is considered as a metabolically less harmful mechanism of WAT growth, than an expansion in volume (hypertrophy) of already existing fat cells which is recognized as a pathological form of WAT remodeling that is associated with adverse cardiometabolic outcomes (1, 2). While hypertrophy in visceral adipose tissue (VAT) is associated with dyslipidemia and a pro-inflammatory state, both subcutaneous white adipose tissue (SAT) and VAT hypertrophy can contribute to systemic insulin resistance (3–8). With progressing obesity, recent studies report a decrease in lipid turnover which can be attributed to elevated rates of triglyceride storage and decreased lipolytic activity (9, 10). Additionally, differences in the secretion pattern of pro- and anti-inflammatory adipokines between small and large fat cells derived from the same individual were reported (11). The similarities and differences regarding measurement methods for adipocyte size and the pathophysiological implications of adipocyte hypertrophy have been extensively and systematically reviewed (12). Large-scale studies trying to unravel the causal transcriptional mechanisms behind adipocyte hypertrophy however, remain scarce (13).

One of the largest public biobanks that includes both tissue histology together with concordant RNA-Seq data is the Genotype-Tissue expression (GTEx) project (14, 15). Since our group previously estimated adipocyte size from over 500 SAT and VAT GTEx samples, we combined adipocyte size and RNA-Seq data from this cohort to investigate the transcriptomic changes that occur between different fat depots and depending on adipocyte hypertrophy (16).

Considering the relevance of fatty acid (FA) composition and fat cell size for WAT function and metabolic disease there is a need to develop novel, less invasive, yet highly accurate methods for the characterization of WAT. However, the feasibility of the simultaneous characterization of both FA composition and adipocyte size by magnetic resonance imaging (MRI) has not been shown yet and remains unclear. Furthermore, the motion sensitivity of magnetic resonance (MR) diffusion-based adipocyte size estimations appears to be challenging in a clinical context given the required long diffusion times (17, 18). Thus, an MR relaxation-based characterization of adipocyte size – which is less sensitive to motion – combined with a chemical shift-based FA decomposition technique is desirable in the clinical context.

Therefore, the present study aimed to provide a comprehensive overview of gene expression changes associated with adipocyte hypertrophy on a variety of representative samples. Further, we investigated two of the potential underlying mechanisms, including mitochondrial activity and FA metabolism. For the latter, adipocyte size was related to FA composition and an advanced multi-parametric MRS-based method for the translational application towards an earlier and non-invasive diagnosis of WAT related metabolic diseases was developed.

## MATERIAL

### Study cohorts, sample origin and conducted experiments

All general study participants’ characteristics, cohorts and related outcome measures are summarized in Table 1.

**Table 1:** Phenotype data, obtained sample types, and measured outcome variables of subjects that were included in the study.

| Cohort | n<br>[male/female] | Age<br>[years] | BMI<br>[kg/m<sup>2</sup>] | Sample types | Outcome variables |
| --- | --- | --- | --- | --- | --- |
| GTEx<br>SAT | 153<br>100/53 | 20 – 29: 9 (5.9 %)<br>30 – 39: 11 (7.2. %)<br>40 – 49: 16 (10.5 %)<br>50 – 59: 46 (30.1 %)<br>60 – 69: 69 (45.1 %)<br>70 – 79: 2 (1.3 %) | N/A | WAT biopsies | RNA-Seq<br>Adipocyte size |
| GTEx<br>VAT | 141<br>92/49 | 20 – 29: 10 (7.1 %)<br>30 – 39: 10 (7.1 %)<br>40 – 49: 16 (11.3 %)<br>50 – 59: 52 (36.9 %)<br>60 – 69: 49 (34.8 %)<br>70 – 79: 4 (2.8 %) | N/A | WAT biopsies | RNA-Seq<br>Adipocyte size |
| GTEx<br>Paired | 99<br>64/35 | 20 – 29: 8 (8.1 %)<br>30 – 39: 8 (8.1 %)<br>40 – 49: 11 (11.1 %)<br>50 – 59: 35 (35.4 %)<br>60 – 69: 35 (35.4 %)<br>70 – 79: 2 (2.0 %) | N/A | WAT biopsies | RNA-Seq<br>Adipocyte size |
| Liposuction | 4<br>0/4 | 39 ± 10 | 27.4 ± 5.8<br>1 N/A | Mature<br>adipocytes<br>separated by<br>size | RNA-Seq<br>Adipocyte size |
| Fatty acids<br>SAT | 22<br>1/21 | 45 ± 12 | 32.6 ± 6.7<br>1 N/A | WAT biopsies | FAME GC-MS<br>Adipocyte size |
| Fatty acids<br>VAT | 12<br>5/7 | 53 ± 15 | 36.0 ± 11.4<br>1 N/A | WAT biopsies | FAME GC-MS<br>Adipocyte size |
| Fatty acids<br>Paired | 7<br>2/5 | 46 ± 12 | 40.7 ± 7.6<br>1 N/A | WAT biopsies | FAME GC-MS<br>Adipocyte size |
| MRS<br>SAT | 16<br>16/0 | 44 ± 12 | 30.6 ± 4.6 | WAT biopsies | MRS-derived FA profile<br>Methylene relaxation |
| MRS<br>VAT | 5<br>3/2 | 51 ± 14 | 34.7 ± 11.4 | WAT biopsies | MRS- derived FA profile<br>Methylene relaxation |

Data is given as means ± SD. Age from GTEx individuals is specified as count (%) in 10 year brackets.

#### GTEx

Parts of the data used in this study to investigate the relationship between adipocyte area and WAT transcriptome originate from the “Genotype-Tissue Expression (GTEx)” project (14, 15). GTEx SAT samples originate from beneath the leg’s skin sample located below the patella on the medial site. VAT was obtained from the greater omentum. Sex and age in 10 year brackets derived from the GTEx publicly available subject phenotypes were used to characterize the donors.

#### Samples for FA composition and MRS measurements

Larger SAT pieces (∼ 500 cm^3^) were obtained from abdominoplasty. Paired SAT and VAT samples were collected during general surgery or abdominal laparoscopic surgery. The described samples were used for MRS measurements, histology-based adipocyte sizing and to determine the FA composition by fatty-acid methyl ester (FAME) GC-MS. Collected phenotypic data included sex, age and BMI.

#### Samples for mature adipocyte isolation

Liposuction material was used to prepare and fractionate mature adipocytes for later isolation of RNA and transcriptomic characterization. Sex, age and BMI were given for the clinical characterization of the samples.

### Cell culture

#### Adipocyte isolation and fractionation

Mature adipocytes were isolated from liposuction material based on collagenase-digestion as described previously (19). After isolation adipocytes were washed with Krebs-Ringer phosphate (KRP) buffer containing 0.1 % BSA for three times. Mature adipocytes were fractionated based on buoyancy in a separating funnel (11, 19). For size separation, 25 ml of mature adipocytes were gently mixed with 50 ml KRP 0.1 % BSA. After 45 s 25 ml were withdrawn from the funnel representing the small adipocyte fraction. The missing volume in the funnel was replaced with KRP 0.1 % BSA and this procedure was repeated 4 times. Similarly, an intermediate fraction was obtained with a flotation time of 20 s. Afterwards, only large cells remained in the funnel which were directly withdrawn.

### Adipocyte size determination

#### Computationally derived adipocyte size estimates from histology samples

GTEx adipocyte area estimates originated from an earlier analysis and published method, the Adipocyte U-Net (16). For additional RNA-Seq analysis of the GTEx samples and to factorize adipocyte area individuals were assigned to one of four bins with equally spaced area intervals (bin_small_, bin_medium_, bin_large_, bin_X-large_). Summary statistics on bin-specific phenotypes and adipocyte areas are given in Table 2.

**Table 2:** Bin and depot specific GTEx phenotypes and adipocyte sizes

|  | SAT |  |  |  | VAT |  |  |  |
| --- | --- | --- | --- | --- | --- | --- | --- | --- |
| **Bin** | ***Small*** | ***Medium*** | ***Large*** | ***X-Large*** | ***Small*** | ***Medium*** | ***Large*** | ***X-Large*** |
| **n** | 20 | 49 | 56 | 28 | 41 | 49 | 40 | 11 |
| **Mean Area**<br>**±**<br>**SD [μm<sup>2</sup>]** | 1,383<br>±<br>241 | 2,261<br>±<br>230 | 3,055<br>±<br>270 | 3,797<br>±<br>235 | 1,089<br>±<br>240 | 2,043<br>±<br>326 | 2,947<br>±<br>255 | 3,860<br>±<br>335 |
| **Area range**<br>**min – max [μm<sup>2</sup>]** | 907 – 1,759 | 1,790 – 2,598 | 2,618 – 3,468 | 3,487 – 4,325 | 482 – 1,503 | 1,556 – 2,540 | 2,570 – 3,403 | 3,580 – 4,612 |
| **Sex (Male/Female)** | 14/6 | 28/21 | 39/17 | 19/9 | 22/19 | 30/19 | 29/11 | 11/0 |
| **Age 20-29 [n/%]** | 3 (15%) | 4 (8.2%) | 2 (3.6%) | 0 (0%) | 8 (20%) | 2 (4.1%) | 0 (0%) | 0 (0%) |
| **Age 30-39 [n/%]** | 2 (10%) | 1 (2.0%) | 7 (12%) | 1 (3.6%) | 4 (9.8%) | 1 (2.0%) | 3 (7.5%) | 2 (18%) |
| **Age 40-49 [n/%]** | 2 (10%) | 6 (12%) | 7 (12%) | 1 (3.6%) | 5 (12%) | 7 (14%) | 4 (10%) | 0 (0%) |
| **Age 50-59 [n/%]** | 7 (35%) | 14 (29%) | 17 (30%) | 8 (29%) | 13 (32%) | 16 (33%) | 17 (42%) | 6 (55%) |
| **Age 60-69 [n/%]** | 6 (30%) | 24 (49%) | 21 (38%) | 18 (64%) | 11 (27%) | 21 (43%) | 14 (35%) | 3 (27%) |
| **Age 70-79 [n/%]** | 0 (0%) | 0 (0%) | 2 (3.6%) | 0 (0%) | 0 (0%) | 2 (4.1%) | 2 (5.0%) | 0 (0%) |

Data is given as means ± SD. Age from GTEx individuals is specified as count (%) in 10 year brackets. Individuals were assigned to one of four bins with equally spaced adipocyte area intervals (small, medium, large, X-large).

#### Histology-based adipocyte sizing of samples used for FA composition measurements and MRS

At least three 5 µm sections with a minimum distance of 100 µm between sections were cut from FFPE WAT. Sections were H&E stained and stitched high definition range bright-field images at 200x magnification were taken (VHX 6000, Keyence). Adipocyte area was determined using the microscope’s in-built image analysis software and diameter was calculated assuming spherical shape (VHX-6000, Version 3.2.0.121). A lower bound threshold of 200 µm^2^ and an upper bound threshold of 16,000 µm^2^ was set to remove artefacts (16). Identified objects were inspected manually and artifacts were excluded from the analysis. Summary statistics on adipocyte area and diameter of the samples that were either used to correlate adipocyte size with FA levels or results derived from MRS are given in Tables S5 & S6, respectively.

#### Mature adipocytes

To measure mature adipocyte diameter a glass-slide was wetted with 200 µl of PBS and approximately 20 µl of adipocytes were pipetted onto the slide. Stitched, bright-field images in high definition range were taken and adipocyte diameter was automatically determined using the microscope’s software (VHX-6000). Identified objects were inspected manually and artifacts were excluded from the analysis.

### RNA isolation, sequencing and differential expression analysis

#### Mature adipocyte RNA isolation

600 µl of adipocytes were pipetted into microtubes containing 600 µl of RLT (Qiagen) 1 % ß- mercaptoethanol (Merck), 200 mg of 0.5 mm ceramic beads (Carl-Roth), inverted and immediately snap-frozen. Samples were allowed to thaw on ice, the RLT 1 % ß-Mercaptoethanol solution was removed and replaced with mirVana lysis buffer (Thermo Fisher Scientific). Samples were lysed for 3x 30 s in a homogenizer filled with dry ice set to 6.0 m/s (FastPrep, MP biomedicals). Samples were centrifuged at 12,000 g for 10 min. at 4 °C. Using an insulin syringe (B. Braun) the adipocyte lysate located underneath the fat layer was transferred to a new tube. All remaining steps were carried out as described by the manufacturer.

#### RNA quality control of mature adipocytes

RNA concentration and 260/280 ratios from mature adipocytes were photometrically determined (Infinite 200 PRO, Tecan). The quality of isolated RNA was assessed based on RNA integrity number (RIN) obtained by using a Bioanalyzer RNA Nano chip (2100 Bioanalyzer, Agilent). Across all samples, high RNA integrity was observed with an average RIN of 9.0 ± 0.8.

#### Mature adipocyte RNA sequencing

Library preparation, sequencing, and reference genome alignment of RNA isolated from mature adipocytes was carried out by the genomics services department of the Broad Institute. Reads were aligned to human genome assembly GRCh37 (hg19) using STAR (20). The number of raw read counts was determined using the feature counts function from the Rsubread package (21). *<u>GTEx RNA-Seq data</u>*

Raw gene read counts and gene transcripts per kilobase million (TPM) were obtained through the GTEx portal from GTEx analysis V8 (dbGap accession phs000424.v8.p2) (14, 15).

#### Differential expression analysis

All RNA-Seq data was analyzed using edgeR (22–24). Multidimensional scaling (MDS) plots and dendrograms were generated to visualize clustering of samples dependent on experimental groups. EdgeR’s quasi-likelihood pipeline was used to test for differential gene expression (24). Differential expression analysis was used to compare gene expression between GTEx-derived SAT and VAT samples. A differential expression model with adipocyte size as a categorical variable (bin_small_ vs. bin_X-large_) was applied to GTEx SAT and VAT samples. Additionally, a model relating gene expression to SAT or VAT adipocyte size expressed as a continuous variable was computed for the GTEx cohort. P-values from GTEx RNA-Seq analysis were FDR corrected and genes with an FDR < 0.05 and a log_2_ fold change > 1 were considered as significant.

In another differential expression analysis, the transcriptome of size-separated adipocytes (small fraction vs. large fraction) was compared. P-values from size-separated adipocytes were FDR corrected and genes with an FDR < 0.05 were considered as significant.

#### Gene set enrichment analysis

Gene set enrichment analysis based on the Kyoto Encyclopedia of Genes and Genomes (KEGG) was carried out using clusterProfiler (25). Data visualization in the form of pathway graphs was achieved using pathview (26).

#### BATLAS analysis

SAT and VAT TPM expression values obtained from the GTEx portal were filtered for the relevant BATLAS marker genes (27). TPM expression values of marker genes that were not expressed in the GTEx cohort were set to zero. Brown and beige adipocyte content estimates per individual and respective WAT depot were obtained from the BATLAS web tools results sheet (https://shiny.hest.ethz.ch/BATLAS/) and mean-centered for all later statistical analysis .

### Adipose tissue fatty acid composition

Adipose tissue FA composition was determined based on FAME GC-MS in samples from abdominoplasty, general surgery or laparoscopy (28). 10 – 20 mg of wet adipose tissue were snap-frozen on dry ice immediately after excision in tubes containing 700 mg lysing matrix D (MP biomedicals). Samples were weighed, allowed to thaw on wet ice, and concentration was set to 0.05 mg/µl using equal parts MeOH and water (LiChrosolv, Merck). Tissue was lysed using a homogenizer (FastPrep) set to 30 s and 6 m/s. Transesterification was carried out as described earlier and FAMEs were extracted using hexane (LiChrosolv, Merck) (28, 29). GC-MS based total FA analysis was performed as previously published (28). Individual FA species were presented as molar percentages of the total FA profile.

### Magnetic resonance spectroscopy

#### Lipid droplet phantom studies

As a validation strategy and to further elucidate the interplay of lipid droplet size and the degree of FA unsaturation in the formation of MR signals, lipid droplet phantoms were manufactured (30). Preceding emulsification, the FA composition and degree of unsaturation of different vegetable oils were validated using FAME GC-MS. Sunflower seed oil and linseed oil were diluted 1:500 and 1:1000 in 3:1 isooctane isopropyl alcohol (LiChrosolv, Merck). Transesterification and GC-MS were carried out similarly to WAT samples as described above. Lipid droplet phantoms were manufactured with the following ratios of sunflower and linseed oil: 1:0, 1:2, 2:1, 0:1. All lipid droplet phantoms were produced with a fat content of 80 % and a water content of 20 % closely resembling human WAT composition (31, 32). For emulsification and conservation purposes the aqueous phase contained 2 % (v/v) tween 80 (Merck) and 0.5 % (m/m) sodium benzoate (Merck). Emulsification was carried out using a colloid mill (Labor-Pilot 2000/4, IKA-Werke) set to 3000, 5000, 8000, and 12,000 rpm to obtain different lipid droplet sizes within the water matrix as described earlier (30, 33, 34). The generated lipid droplet size spectrum was determined using a laser diffraction particle size analyzer ((Mastersizer 2000 with Hydro 2000S dispersing unit, Malvern Instruments).

#### Adipose tissue sample preparation

All human WAT samples originating from abdominoplasty, general surgery or laparoscopy used for MRS were fixed in 4 % formaldehyde (Histofix, Carl-Roth) for 24 hours and immediately measured afterwards.

#### Magnetic resonance spectroscopy

Lipid droplet phantoms and WAT were measured using a single-voxel short-TR multi-TI multi-TE (SHORTIE) stimulated echo acquisition mode (STEAM) MRS acquisition scheme (Figure 7A) (35). Both lipid droplet phantoms and WAT samples were measured using equivalent sequence parameters: Inversion time (TI) of 8, 83, 233, 458, 833, 1133 ms, echo time(TE) of 10, 15, 20, 25, 70 ms, mixing time (TM) of 16 ms, minimum repetition time (TR) of 801 ms, τ of 774 ms, 4 phase cycles in 4 averages, 2048 sampling points, spectral bandwidth of 3000 Hz, default voxel-size of 12x12x12 mm^3^ and a total scan time of 02:32 min. Spectra were obtained for all combinations of TI and TE. All measurements were performed at room temperature (21±1°C) on a clinical whole-body 3T MRI scanner (Ingenia Elition X, Philips Healthcare, The Netherlands). For signal reception, either the clinical 8-channel small extremity coil or the clinical 8-channel wrist coil was used depending on sample size. The acquired MRS data was processed using SVD-based coil combination followed by simple averaging and zero-order phase correction (36). Signal fitting was carried out using a time domain-based joint-series model fitting routine implemented in MATLAB (R2019b) using NL2SOL (37). The SHORTIE signal as a function of time S(t) was modeled as:

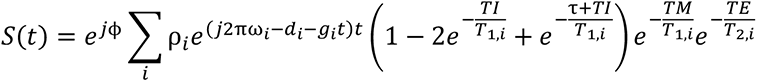

where *ϕ* represents a common phase term, *ρ_i_* is the proton density, *ω_i_* is the precession frequency, *d_i_* and *g_i_* are the Lorentzian and Gaussian damping factors, and T_1,i_ and T_2,i_ are the relaxation times of the *i*th frequency component, respectively. TI is the inversion time; TM is the mixing time and TE is the echo time. The proton MR-based FA profile characterization is based on the assumption that all MR-detectable FAs are present in the form of triglycerides and that all MR-detectable signals originate from water and triglycerides. Consequently, a 11-peak-signal-model was fitted with in total 21 degrees of freedom including constraints for a 10-peak-triglyceride-model and relaxation properties (Table S7) (38). In course of the parametrization of the triglyceride model, the FA profile characterization results from three triglyceride characteristics: the mean number of double bounds per triglyceride (ndb), the mean number of methylene interrupted double bounds per triglyceride (nmidb) and the mean fatty acid carbon chain length (CL).

### Statistical analysis

Data analysis was carried out using R (39). If not specified otherwise all results were given as mean ± SD. According to the sample distribution, variance, and experimental setting (paired vs. unpaired) parametric tests (paired samples t-test, independent samples t-test) or non-parametric tests (Mann-Whitney test, Wilcoxon signed-rank test) were used to test for/against differences between groups. Similarly, Pearson or Spearman correlation analysis was applied to investigate the association between continuous variables. One-sided tests were used within the BATLAS analysis to investigate whether a decrease in brown/beige adipocyte content was observed with larger fat cell size. Bonferroni correction was used to adjust for multiple testing in the FA composition analysis. Across all analyses p-values < 0.05 were considered significant. Linear regression analysis of the MRS data was carried out in python (V3.6.10) using the scipy package (V1.4.1).

### Study approval

All tissue donors gave written informed consent. The study protocols were approved by the ethics committee of the Technical University of Munich (Study №: 5716/13, 1946/07, 409/16s)

## RESULTS

### SAT and VAT display different gene expression signatures

To assess the overall gene expression differences between SAT and VAT, GTEx derived bulk tissue RNA-Seq data was used. As visible by two distinct clusters in the MDS plot and two separate branches in the dendrogram, global SAT and VAT gene expression were clearly distinguishable from each other (n_paired_ = 99, Figure 1A, 1B). Differential gene expression analysis revealed a total of 689 genes characterized by positive fold changes (FC) and thus higher expression levels in SAT compared to VAT (FDR < 0.05, log_2_ (FC) >1; Figure 1C, Table S1_A), while 1437 genes were higher expressed in VAT compared to SAT (FDR < 0.05, log_2_ (FC) >1; Figure 1C, Table S1_A). A large fraction of the top differentially expressed targets were identified as genes of “developmental origin” (*HOX family, IRX3, IRX5, TBX15, SHOX2, NR2F1;* Figure 1C). Amongst genes that were not related to cell fate and development, *MMP3*, known to be involved in the breakdown of extracellular matrix, showed strong enrichment in SAT (log_2_(FC) = -7.82, FDR = 2.71*10^-31^ Figure 1C). On the contrary, *ALOX15* expression was multiple fold higher in VAT compared to SAT and among the top 20 differentially expressed genes in the present analysis (log_2_(FC) = -10.11, FDR = 2.49*10^-62^ Figure 1C, Table S1_A). ALOX15, is involved in the metabolism of poly-unsaturated FAs (PUFAS), producing both pro- and anti-inflammatory mediators, dependent on its FA substrates (40). KEGG gene set enrichment analysis revealed differences between the two different depots revolving around chemokine and cytokine related signaling (*hsa04060*:Cytokine-cytokine receptor interaction, FDR = 1.63e^-06^; *hsa04062*:Chemokine signaling pathway, FDR = 3.44e^-04^), inflammatory processes (*hsa04668*:TNF signaling, FDR = 0.0084; *hsa04064*:NF-kappa b signaling, FDR = 7.51e^-04^) as well as energy metabolism (*hsa00190*:Oxidative phosphorylation, FDR = 0.010). An overview of the top 20 enriched KEGG pathways with an adjusted p-value < 0.05 is given in Figure 1D and all significantly enriched KEGG pathways are depicted in Table S1_B.

**Figure 1:**
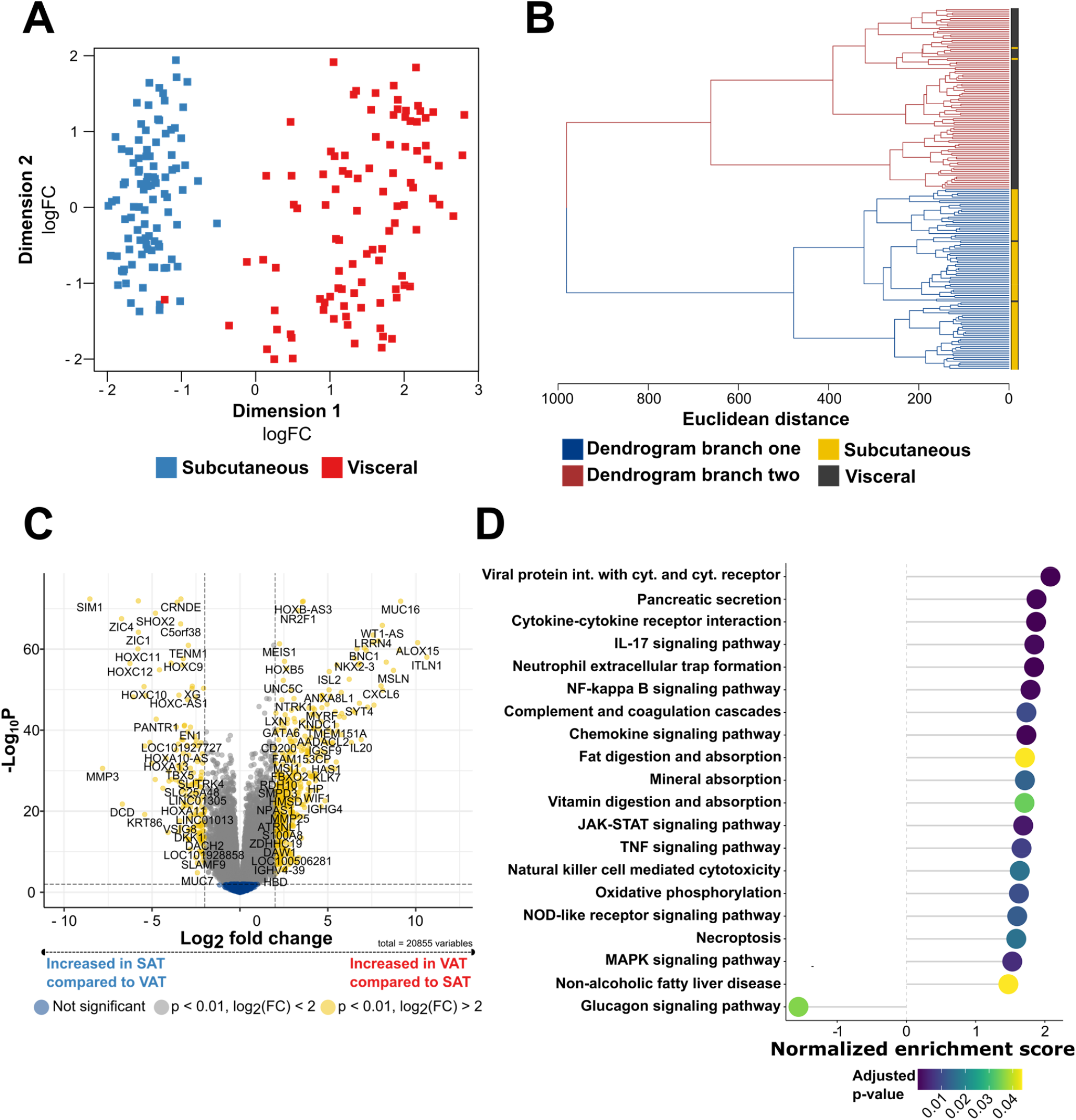
SAT and VAT are characterized by different transcriptomic signatures. **(A)** MDS plot displaying the first two dimensions of the data. **(B)** Hierarchical clustering of SAT and VAT samples based on euclidean sample distances. **(C)** Volcano plot showing differentially expressed genes between SAT and VAT **(D)** Lollipop plot displaying the top 20 enriched KEGG pathways with an adjusted p-value < 0.05 from gene set enrichment analysis, thereby comparing the expression of KEGG pathway associated genes from SAT and VAT depots.

### The SAT and VAT transcriptome is altered with adipocyte hypertrophy

To evaluate the link between adipocyte area and gene expression profiles, individuals from GTEx were stratified into four equally spaced adipocyte area bins (16). A 2.75 fold change in SAT mean adipocyte area was observed between individuals assigned to either bin_small_ (n = 20, mean adipocyte area = 1,383 ± 241 µm^2^) or bin_X-large_ (n = 28, mean adipocyte area = 3,797 ± 235 µm^2^) (Table 2). Substantial changes in SAT gene expression were observed when comparing small with X-large adipocytes, including increased expression of 621 genes in bin_X-large_ compared to bin_small_ (FDR < 0.05, log_2_(FC) > 1; Figure 2A, Table S2_A). 369 genes were characterized by higher expression in the small compared to the X-large bin (FDR < 0.05, log_2_(FC) > 1; Table S2_A). Amongst the top-ranked genes, *SLC27A2* exhibited significantly higher expression in the group with small adipocytes (FDR = 6.33e^-09^, log_2_(FC) = -3.90, Figure 2A, Table S2_A). Further *SLC27A2* ranked highest by FDR in a differential expression analysis where SAT adipocyte area was set as a continuous covariate (n = 153, FDR = 1.74e^-14^; Figure 2C, Table S2_B). SLC27A1 (alias FATP1), the very long-chain FA transporter that is mainly expressed in WAT and that has been previously associated with obesity was not found to be differentially expressed concerning SAT adipocyte area (FDR_binned_ = 0.70, FDR_continuous_ = 0.33) (41, 42). Furthermore, *EGFL6* which has been previously related to obesity and the proliferation of human adipose mesenchymal stem cells was found to be higher expressed in individuals with larger SAT adipocyte areas (FDR = 2.83e^-05^, log_2_(FC) = 4.67, Figure 2A)(43). In the SAT differential gene expression model with adipocyte area as a continuous covariate *SLC27A2* (FDR = 1.74e^-14^), *AMN* (5.89e^-14^), *TC36* (FDR = 8.02e^-14^)*, PTPN3* (FDR = 6.03e^-13^), and *TOX3* (2.46e^-12^) were found as the top 5 genes by FDR (Figure 2C, top row; Table S2_B). An overview of all genes with an FDR < 0.05 overlapping between the binned and continuous model is given in Table S2_C and Figure S1A. The present work is consistent and expands on recent studies to link gene expression with obesity, adipogenesis and fat cell size (12, 44). Amongst others, negative associations between SAT adipocyte area, *SLC2A4* (alias *GLUT4*, FDR =1.72e^-04^), *INSR* (FDR = 0.0013) and *CS* (FDR = 0.0016) were found, while *TNF* (3.07e^-04^) and *EGFL6* (0.014) were positively related with adipocyte area (Figure 2C, bottom row; Table S2_B). KEGG gene set analysis in SAT revealed significant positive enrichment of genes involved in immune-related and inflammatory processes for individuals assigned to the bin_XL_ group (*hsa04640*:Hematopoietic cell lineage, FDR =1.05e^-07^; *hsa04060*:Cytokine-cytokine receptor interaction, FDR = 5.47e^-09^; *hsa04064*:NF-kappa B signaling pathway, FDR = 6.48e^-08^; Figure 2B; Table S2_D). On the contrary, negative enrichment was seen for gene sets involved in key adipose tissue metabolic processes, such as *hsa04923*:Regulation of lipolysis in adipocytes (FDR = 1.13e^-05^; Figure 2B; Table S2_D) and *hsa00620*:Pyruvate metabolism (FDR = 6.48e^-06^; Figure 2B; Table S2_D). The analysis further revealed highly significant negative enrichment for *hsa03010*:Ribosome (FDR = 5.47e^-09^, Figure 2B; Table S2_D) and *hsa00190*:Oxidative phosphorylation (FDR = 5.47e^-09^, Figure 2B; Table S2_D). Pathway analysis confirmed this finding and revealed a consistent negative enrichment of subunits from the mitochondrial respiratory chain in SAT from individuals with enlarged adipocytes (Figure 2D).

**Figure 2:**
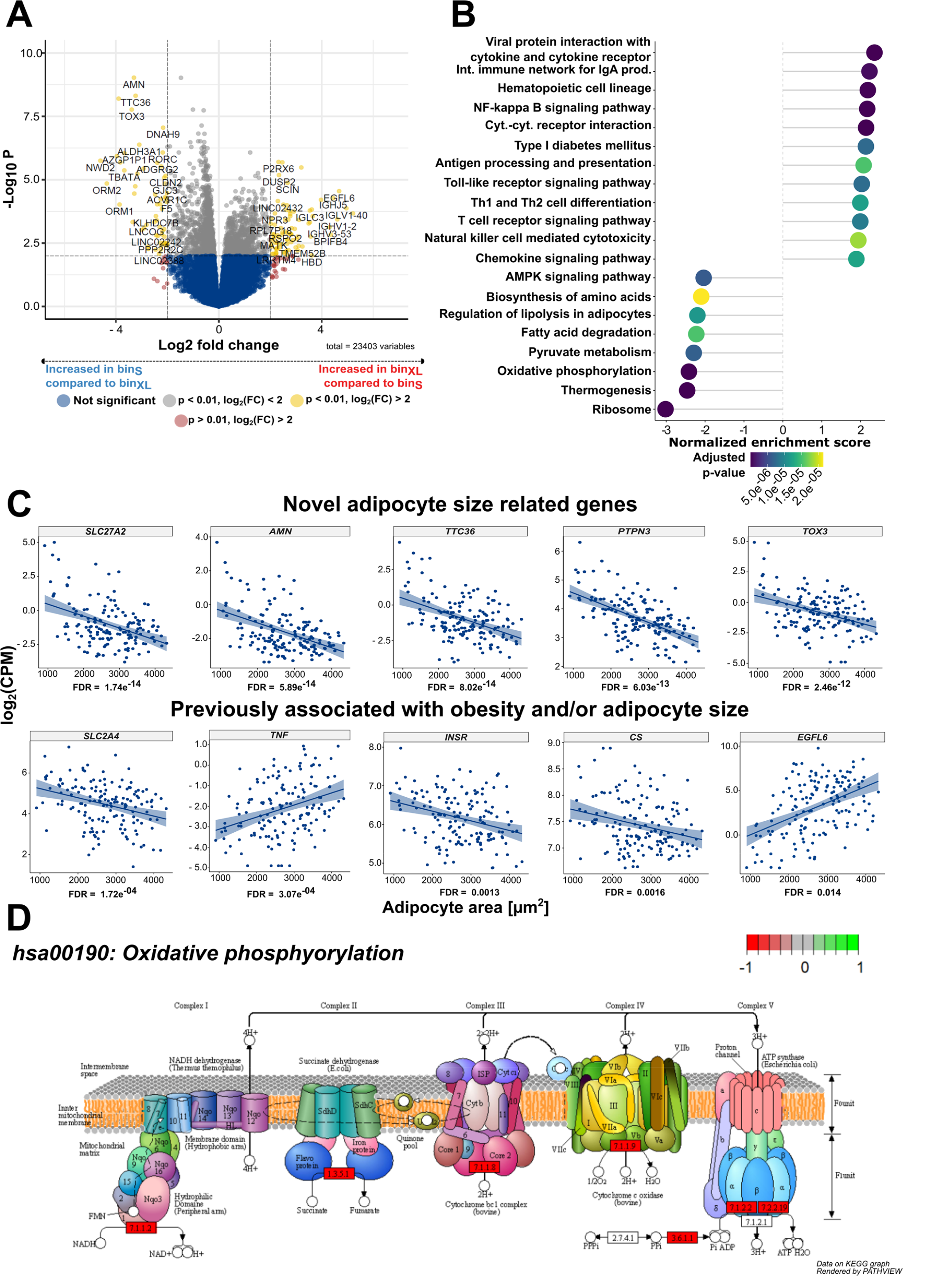
Adipocyte area is related to gene expression in SAT. **(A)** Volcano plot showing DE of genes between the bin_small_ (n = 20) and bin_X-Large_ (n = 28) in SAT. **(B)** Lollipop plot displaying the top 20 enriched KEGG pathways with an adjusted p-value < 0.05 from gene set enrichment analysis in SAT. The “Human Diseases” category of the KEGG pathway database was not displayed on the graph. **(C)** Scatterplots displaying the results of the continuous model-based analysis that was applied to GTEx RNA-Seq data from tSAT. Log2 counts per million (CPM) are plotted against adipocyte area for 153 individuals. FDR is given for each gene below the plot. The upper half of the figure shows 5 of the top-ranked genes from the continuous model-based RNA-Seq analysis *(SLC27A2, AMN, TTC36, PTPN3, TOX3).* In the lower half 5 exemplary genes that are already known to be related to WAT biology and fat cell size that were also significant (FDR < 0.05) in our continuous model are depicted *(SLC2A4, TNF, INSR, CS, EGFL6).* A regression line including the standard error in a lighter shade of blue was added to the plots to visualize the relationship between gene expression and adipocyte area. (D) KEGG Pathview graph depicting the different complexes (I-V) of the respiratory chain (*hsa00190*: Oxidative phosphorylation). Red (-1) indicates mitochondrial complexes that are underrepresented in SAT from the bin_X-Large_ group compared to the bin_small_ group. In contrast, green (+1) displays complexes from the electron transport chain that are overrepresented in SAT from the bin_X-Large_ group in relation to the bin_small_ samples.

VAT adipocyte area binning resulted in a 3.54 fold change in fat cell size between the smallest and largest bin (Table 2). Differences in VAT adipocyte area were reflected by size-bin specific transcriptomic signatures with 500 genes showing higher expression in bin_X-large_ compared to bin_small_ and 327 genes showing higher expression in bin_small_ versus bin_X-Large_ (FDR < 0.05, log_2_(FC) > 1 or log_2_(FC) < -1, respectively; Table S3_A). A differential gene expression model with adipocyte area as a continuous covariate was applied to account for unequal group sizes and sex distributions. The continuous model revealed a higher number of significant genes with an FDR < 0.05 (continuous: 4,194 genes; categorical: 1,896 genes; Figure S1B, Table S3_B) most likely due to higher sample sizes (n_cont_ = 141). Differentially expressed genes from the categorical model were overlapping with the genes from the continuous model with only 208 differentially expressed genes specific to the categorical model (Figure S1B, Table S3_C). An overview of ten exemplary genes that were found amongst the top 20 adipocyte size dependent genes in VAT by the continuous model is given in Figure 3A. All differentially expressed genes from the continuous model are listed in Table S3_B. With both models a strong negative relationship between *ELOVL6* and adipocyte area was found (FDR_continuous_ = 1.92e^-12^, FDR_cat_ = 9.88e^-05^, log_2_(FC)_cat_ = -4.21, Figure 3A). As depicted in Figure 3B, KEGG gene set enrichment analysis revealed significant negative enrichment for genes involved in metabolic pathways (*hsa00620*:Pyruvate metabolism, FDR = 3.66e^-05^ *hsa00020*:Citric acid cycle (TCA), FDR = 1.17e^-05^ *hsa01212*:Fatty acid metabolism, FDR = 9.70e^-07^ Table S3_D), signaling (*hsa03320*:PPAR signaling pathways, FDR = 8.80e^-06^; *hsa04910*:Insulin signaling, FDR = 4.21e^-05^; Table S3_D) and energy homeostasis (*hsa00190*:Oxidative phosphorylation, FDR = 2.29e^-05^; *hsa04714*:Thermogenesis, FDR = 6.60e^-09^; Table S3_D). In line with findings from KEGG enrichment analysis suggesting that the FA metabolism of individuals with large adipocytes is perturbed and an inverse relationship between adipocyte area and *ELOVL6*, a negative relationship between adipocyte area and *FASN* expression (FDR_Cat_ = 0.004, log_2_(FC)_Cat_ = -3.23; FDR_Cont_ = 2.02e^-06^) was observed in both models (Figure 3C). In agreement with the KEGG pathway analysis, the categorical and the continuous model revealed reduced expression of *UCP-1* in VAT from individuals with enlarged fat cells (FDR_Cat_ = 0.017, log_2_(FC)_Cat_ = -4.74; FDR_Cont_ = 1.57e^-05^ Figure 3C). A detailed visualization of the KEGG thermogenesis pathway indicated concordant negative enrichment of genes involved in lipolysis (*ATGL*, *HSL*, *PLIN*), respiration and energy dissipation (*UCP-1*) in VAT from individuals with hypertrophic adipocytes (Figure 3D). Together, adipocyte hypertrophy in both SAT and VAT was associated with decreased marker gene expression of genes involved in processes crucial for WAT function and homeostasis, i.e. FA metabolism and mitochondrial activity.

**Figure 3:**
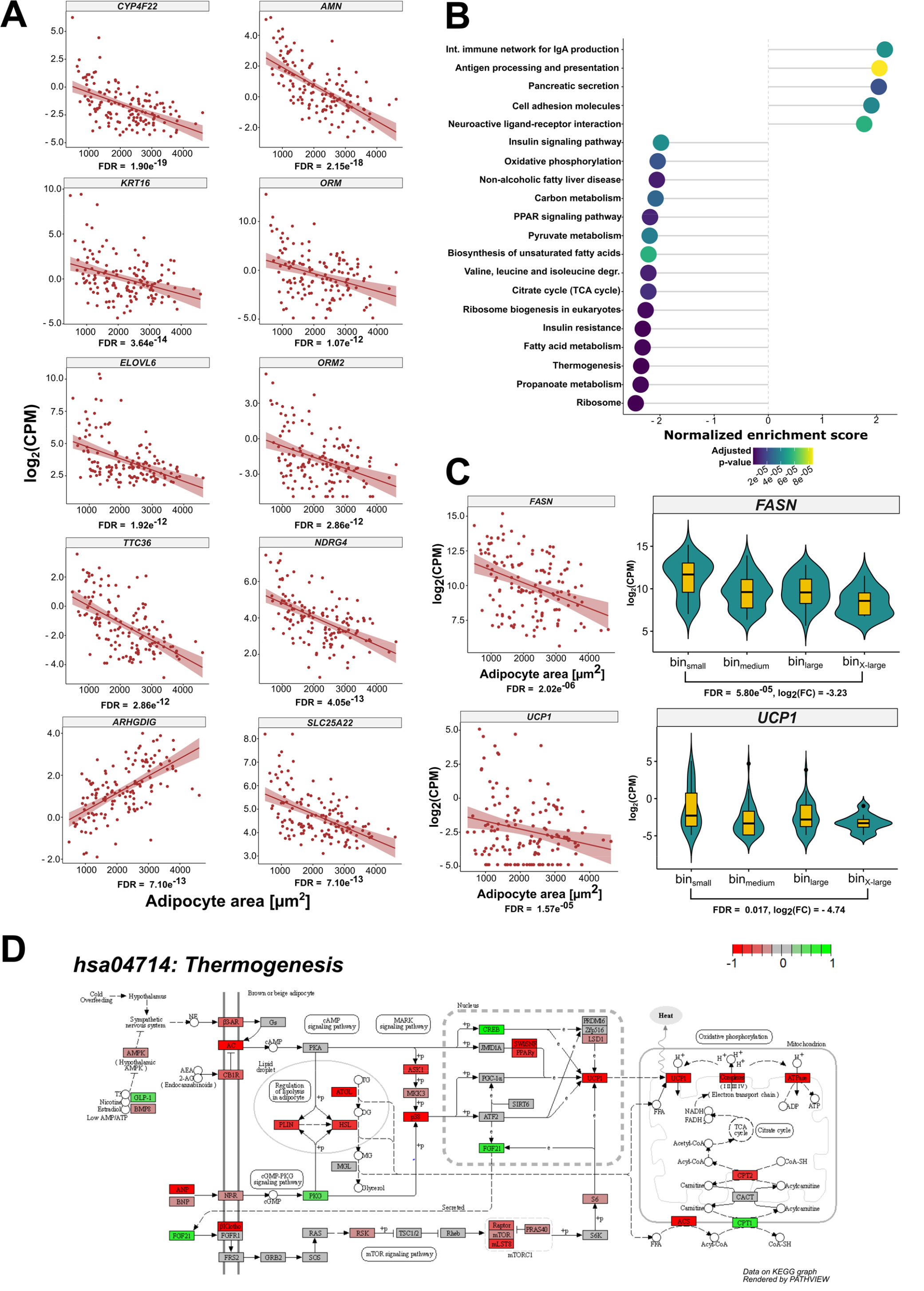
Adipocyte area is related to gene expression in VAT. (A) Scatterplots displaying the results of the continuous model-based analysis that was applied to GTEx RNA-Seq data from VAT. Log2 counts per million (CPM) are plotted against adipocyte area for 141 individuals. FDR is given for each gene below the plot. Displayed is the relationship between adipocyte area and gene expression for 10 exemplary genes that were ranked highest in the analysis according to their FDR. A regression line including the standard error in a lighter shade of red was added to the plots to visualize the relationship between gene expression and adipocyte area.(B) Lollipop plot displaying the top 20 enriched KEGG pathways with an adjusted p-value < 0.05 from gene set enrichment analysis in VAT. The “Human Diseases” category of the KEGG PATHWAY database is not displayed on the graph. (C) Scatterplot and violin plot assessing the relationship between the expression of *FASN*, *UCP1* and adipocyte area. (D) KEGG Pathview graph depicting the different complexes and genes involved in the canonical thermogenesis pathway (hsa04714:Thermogenesis). Red (-1) indicates thermogenesis related genes that are underrepresented in VAT from the bin_X-Large_ group compared to the bin_small_ group. In contrast, green (+1) displays thermogenesis genes that are overrepresented in VAT from the bin_X-Large_ group in relation to the bin_small_ samples. Genes depicted in grey (0) did not show differences between the two groups.

### Adipocyte size is inversely associated with the thermogenic adipocyte content of SAT and VAT

To further elucidate the link between adipocyte area and mitochondrial activity, the brown adipose tissue atlas (BATLAS) deconvolution tool was used to estimate brown and beige adipocyte content in GTEx WAT biopsies as a proxy of thermogenesis and browning capacity (27). BATLAS analysis revealed substantial differences in estimated thermogenic adipocyte content dependent on WAT depot and adipocyte area (Table 3). A 6.13 % higher brown/beige adipocyte content was predicted in paired VAT compared to SAT derived from 99 individuals (p_paired t-test_ = 2.07e^-06^ Figure 4A, Table 3). In both VAT and SAT, BATLAS predicted a decrease in brown/beige adipocyte content with hypertrophic (bin_X-Large_) compared to small adipocytes (bin_small_). A 8.79 % and 13.57 % reduction in brown/beige adipocyte content was observed for SAT and VAT, respectively (SC: n_binSmall_ = 20, n_binX-Large_ = 28, p_Wilcoxon_ = 0.0013; VC: n_binSmall_ = 41, n_binX-Large_ = 11, p_t-test_ = 0.0038; Figure 4B, 4C, Table 3). Findings from the binned analysis were confirmed by correlating the estimated brown adipose tissue (BAT) content with adipocyte area across all samples from both adipose depots. In SAT, a negative relationship between brown/beige adipocyte content and adipocyte area was found (n = 153, ρ_spearman_ = -0.16, p = 0.026, Figure 4B). Similar results were observed in VAT where adipocyte area was inversely associated with BAT content estimated by BATLAS (n = 141, r_Pearson_

**Figure 4:**
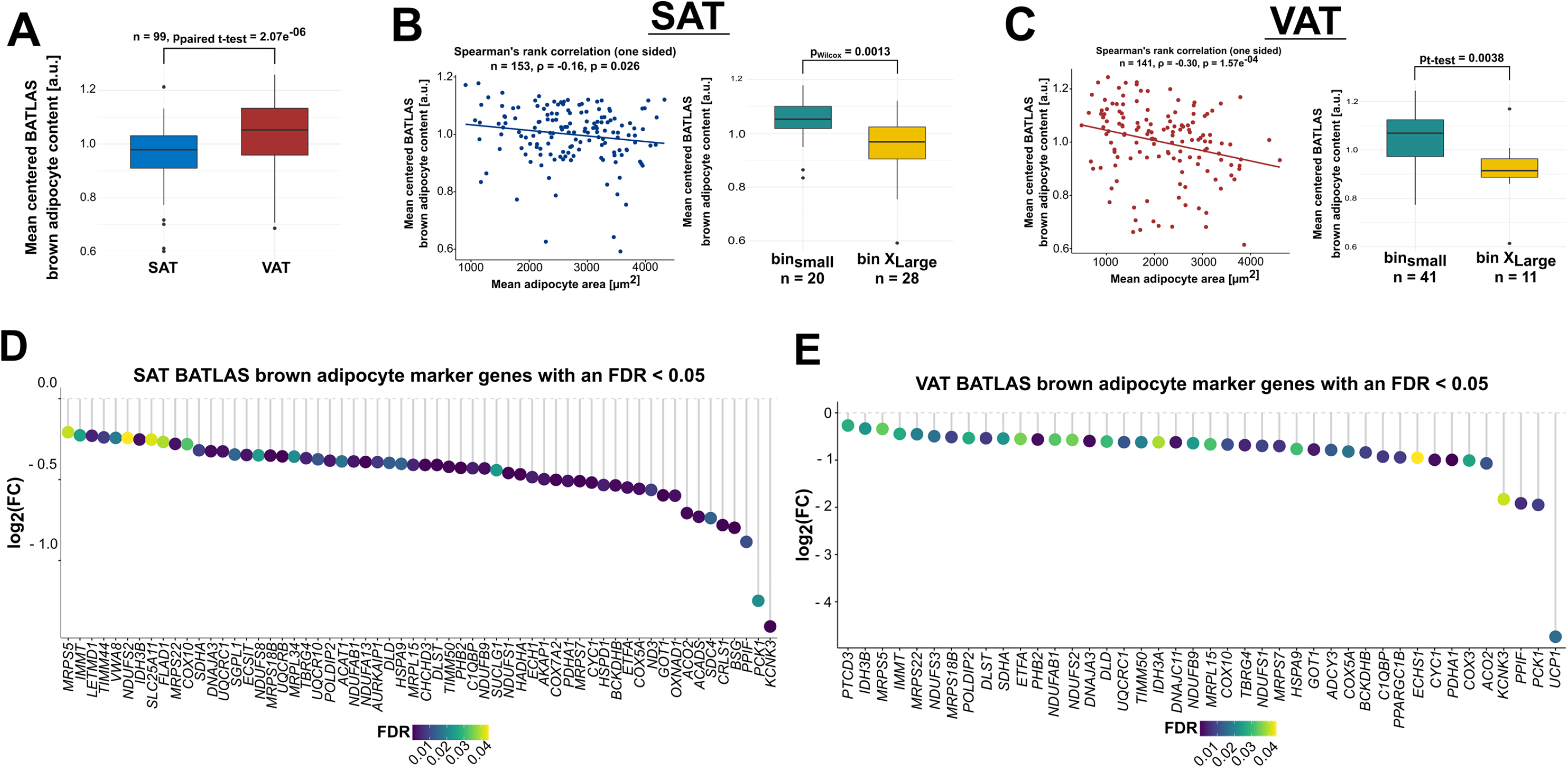
BATLAS analysis indicates differences in brown adipocyte content with adipocyte hypertrophy. **(A)** Tabular overview from BATLAS analysis for the depot and size bin-specific changes in estimated brown adipocyte content. Additionally, the number of the brown (n_brown marker genes_ = 98) and white (n_white marker genes_ = 21) marker genes from BATLAS that were differentially expressed with an FDR < 0.05 for the relevant group-wise comparisons are listed. **(B)** Boxplot comparing the mean-centered BATLAS estimated brown adipocyte content of paired SAT and VAT biopsies. **(C)** Scatterplot depicting the correlation between estimated brown adipocyte content and adipocyte area in SAT (left). Boxplot comparing the estimated brown adipocyte content between individuals with small (bin_small_) and large subcutaneous adipocytes (bin_X-large_). **(D)** Scatterplot depicting the correlation between estimated brown adipocyte content and adipoyte area in VAT (left). Boxplot comparing the estimated brown adipocyte content between individuals with small (bin_small_) and large visceral adipocytes (bin_X-large_) (right). **(E)** Lollipop plot depicting all BATLAS brown adipocyte marker genes that were found to be differentially expressed with a FDR < 0.05 in SAT group-wise comparisons between the bin_small_ and bin_X-Large_ group. **(F)** Lollipop plot depicting all BATLAS brown adipocyte marker genes that were found to be differentially expressed with a FDR < 0.05 in VAT group-wise comparisons between the bin_small_ and bin_X-Large_ group.

**Table 3:** Tabular overview from the BATLAS analysis for the depot and size bin-specific changes in estimated brown adipocyte content as well as brown and white marker genes.

|  | % Change in brown adipocyte content predicted by BATLAS | BATLAS BAT marker genes with a FDR < 0.05 | BATLAS WAT marker genes with a FDR < 0.05 |
| --- | --- | --- | --- |
| SAT vs VAT | + 6.13 % | 5 ↓ / 61 ↑ | 9 ↓ / 5 ↑ |
| SAT<br>Bin<sub>small</sub> vs- bin<sub>X-Large</sub> | - 8.79 % | 60 ↓ / 0 ↑ | 6 ↓ / 2 ↑ |
| VAT<br>Bin<sub>small</sub> vs- bin<sub>X-Large</sub> | - 13.57 % | 42 ↓ / 0 ↑ | 1 ↓ / 1 ↑ |

= -0.3, p = 1.57e^-04^, Figure 4C). A slightly higher overall BAT content and a stronger negative relationship with adipocyte area were seen for VAT. In both depots, multiple of the 98 BAT marker genes published by *Perdikari et al.* (27) were also found to be significant with an FDR < 0.05 between the bin_small_ and bin_X-Large_ groups (Figure 4D, 4E). Among the significant genes, greater fold changes of BAT marker genes were observed in VAT compared to SAT (Figure 4D, 4E). From all BATLAS BAT marker genes *KCNK3* showed the strongest downregulation in SAT from individuals with large adipocytes (Figure 4D; log_2_(FC) = -1.41, p = 2-54e^-04^). In VAT, *UCP-1* showed the strongest downregulation in individuals with large adipocytes (Figure 4E; log_2_(FC) = -4.74, p =0.017). Similarly, the continuous model indicated a significant negative relationship between *UCP-1* expression and VAT adipocyte area (FDR = 1.57e^-05^).

### RNA-Seq reveals differences in the transcriptomic signature between small and large adipocytes from the same individual involving FA metabolism and inflammation

Results concerning the investigations on bulk adipose tissue from individuals with diverse genetic backgrounds (GTEx cohort) suggest substantial differences in transcriptomic signatures between individuals with large and small fat cells. To clarify, whether the observed differences in tissue can be explained by the transcriptomic signatures of large adipocytes themselves an investigation was carried out on the intra-individual gene expression patterns of size-separated mature adipocytes. Size separation based on buoyancy resulted in an average size difference of 2.98 fold between the small and large fraction (Figure 5A). RNA-Seq revealed 583 genes to be differentially expressed (FDR < 0.05) between small and large adipocytes (Table S4_A). 480 genes were upregulated in the large fraction, while 103 genes were higher expressed in the small fraction (Figure 5B; Table S4_A). KEGG pathway analysis revealed pathways analogous to the cross-sectional GTEx cohort to be enriched in the large (*hsa04151*:PI3K-Akt signaling, FDR = 0.014; *hsa03010*:Ribosome, FDR = 0.0049) and small fraction (*hsa04512*:Carbon metabolism, FDR = 0.0013; *hsa00020*:Citrate cycle, FDR = 0.011; *hsa00620*:Pyruvate metabolism, FDR = 0.015), respectively (Figure 5C; Table S4_B). In contrast to the GTEx tissue-based analysis, OXPHOS and thermogenesis were not significantly enriched in the KEGG gene set enrichment analysis. Further, no major differences in BATLAS predicted thermogenic adipocyte content was observed (data not shown). In conclusion, intra-individual variance of adipocyte size did not alter the expression of genes related to OXPHOS and thermogenesis. The evidence for a decreased expression of genes related to respiratory chain function and thermogenesis was therefore limited to the GTEx RNA-Seq analysis assessing inter-individual differences in gene expression with regard to adipocyte area.

**Figure 5:**
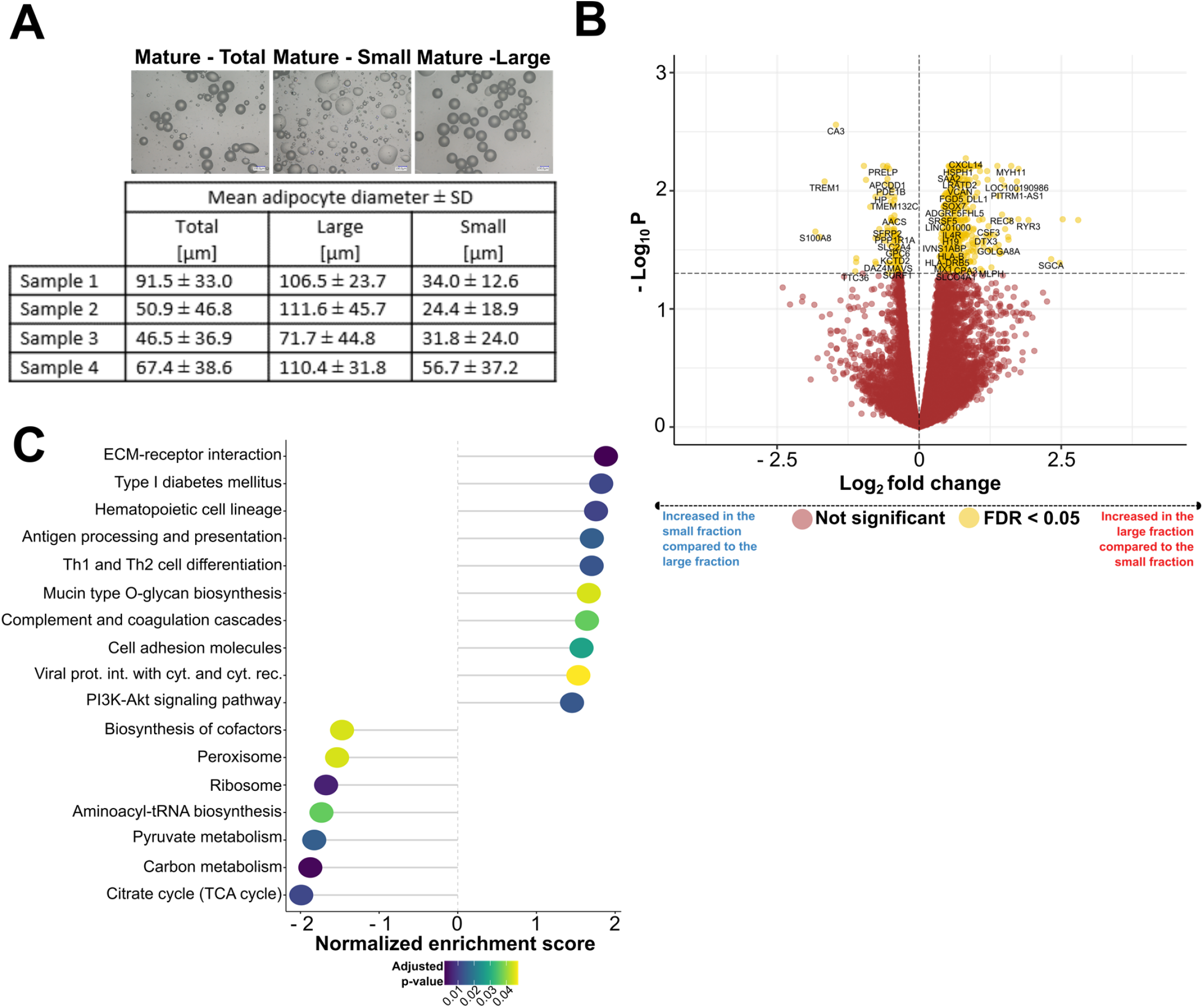
The transcriptomic signatures of size-separated mature adipocytes. **(A)** Results from mature adipocyte size-separation experiments. Representative images of the total, small and large adipocyte fraction from one donor are shown in the top half of the graph. A tabular overview listing the mean adipocyte diameter ± SD per fraction across all four different donors is given in the bottom half. **(B)** Volcano plot depicting the DE of genes between the small and large adipocyte fraction. **(C)** Lollipop plot displaying the top 20 enriched KEGG pathways with an adjusted p-value < 0.05 from gene set enrichment analysis comparing large and small fat cells that were size-separated based on buoyancy. The “Human Diseases” category of the KEGG PATHWAY database is not displayed on the graph.

### Adipose tissue FA composition is related to adipocyte size

To further understand the link between adipocyte area and FA metabolism that the data revealed in the gene set enrichment analysis, i.e. size-, and depot-dependent expression of de novo lipogenesis (DNL) and FA elongation related marker genes, the relationship between WAT FA composition and adipocyte diameter was investigated. Comparisons of the FA composition between SAT and VAT revealed an increased relative abundance of the saturated FA species lauric acid (12:0, p = 0.05), myristic acid (14:0, p = 0.026) and arachidic acid (20:0, p = 0.0091) in VAT (Figure 6A). In contrast, ω-3 and ω-6 PUFAs with chain lengths of 20 or 22 carbon atoms were significantly higher in SAT (Figure 6A). Out of all FA species the ω-6 FA, arachidonic acid (ARA) showed the highest significance (FA20:4 (n-6), p = 0.00082, Figure 6A). Similar to depot-specific FA patterns the results indicate that FA composition is dependent on adipocyte diameter in both depots. In SAT a positive relationship between C20 & C22 PUFAs and adipocyte diameter was observed, while an inverse association was revealed predominantly for medium-chain saturated FAs in VAT (Figure 6B, 6C). Associations between SAT adipocyte diameter and capric acid (FA 10:0, r_Pearson_ = -0.71, p = 2.28e^-04^), eicosapentaenoic acid (FA20:5 (n-3); r_Pearson_ = 0.70, p = 2.66e^-04^) and ARA ( FA20:4 (n-6); r_Pearson_ = 0.70, p = 1.29e^-03^) were observed (Figure 6B, 6C). Inverse relationships of VAT adipocyte diameter with myristic acid (FA14:0, r_pearson_ = -0.88, p = 1.51e^-04^), pentadecylic acid (FA15:0, r_pearson_ = -0.84, p = 5.58e^-04^) and arachidic acid (FA20:0, r_pearson_ = -0.71, p = 0.01) were detected (Figure 6B, 6C).

**Figure 6:**
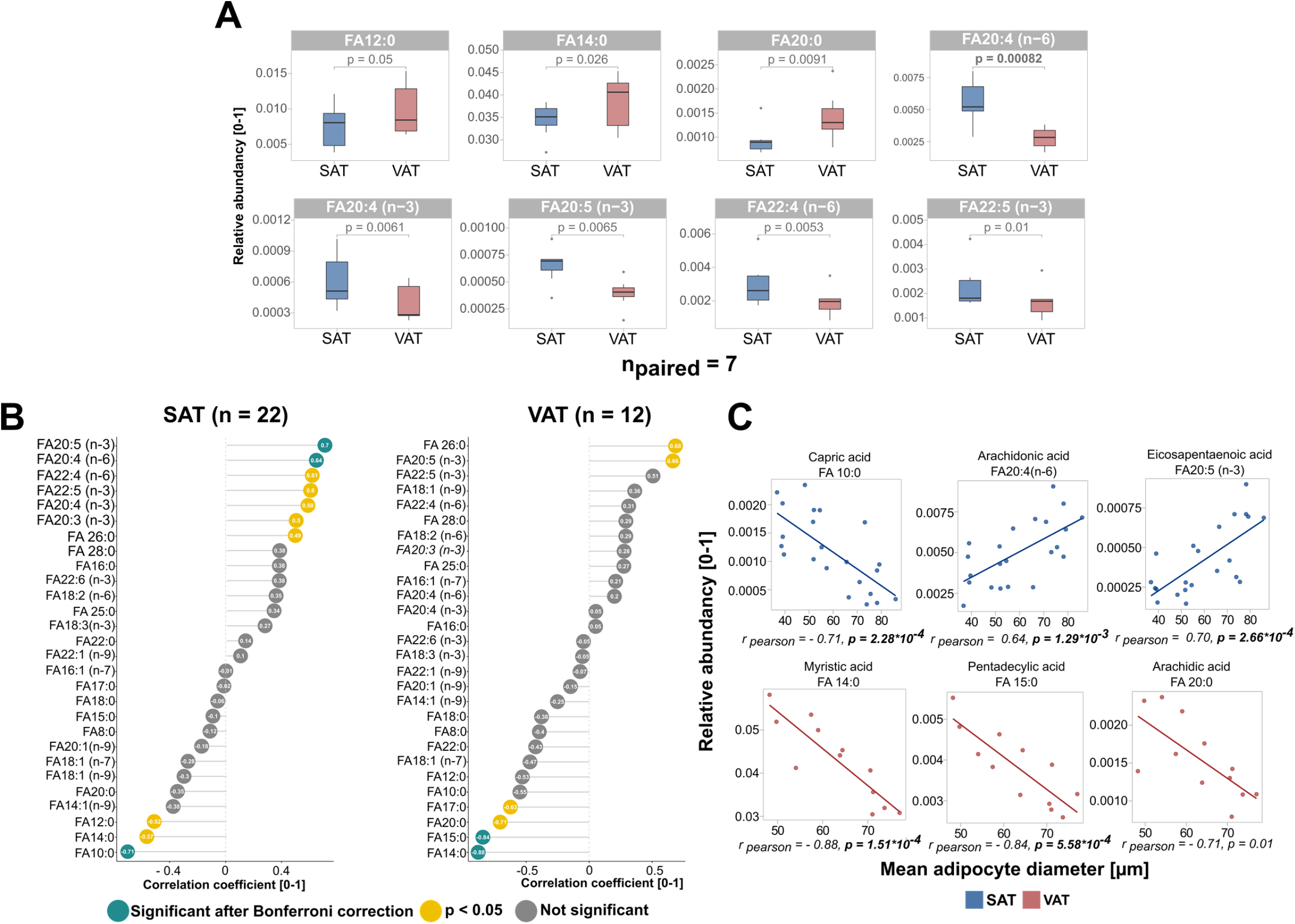
Differences in fatty acid composition with regard to adipose depot and fat cell size. **(A)** Boxplots showing FA species that were found to show different abundancies in paired, SAT, and VAT. SAT samples are colored in red, while VATsamples are shown in blue. Out of the 28 FA species that were measured all FAs with a p-value < 0.05 are displayed. P-values written in bold are the ones that remained significant after Bonferroni adjustment for multiple testing (p_bonf_ = 0.05/28 = 0.0018). **(B)** Lollipop plot showing the correlation between adipocyte diameter and all 28 FA species assessed. SAT samples are displayed on the left, while VAT samples are shown on the right. **(C)** Scatterplots displaying the three FA species showing the strongest association with mean adipocyte diameter in SAT (blue) and VAT (red).

### MRS is suitable for the non-invasive and in-parallel characterization WAT fat cell size and FA composition

To facilitate the translation of adipocyte size associated changes in FA and transcriptomic signatures into clinical practice an MRS-based technique for the simultaneous characterization of FA composition and adipocyte cell size-based relaxation properties was developed. As previously shown by our group, water-fat emulsions are a valuable model system to generate lipid droplets with highly defined size ranges for the MR-based morphological characterization of lipid-rich samples (water/fat content & droplet size) (18, 30).

Independent of the ratio of sunflower to linseed oil, increasing lipid droplet diameters were observed with decreasing revolutions per minute of the colloid mill (Figure S2 A – F, Table S8). At similar rpms a trend towards smaller lipid diameter with increasing linseed oil content was observed (Table S8). As published, linoleic acid (18:2 n-6) was the most abundant FA in sunflower oil (Figure S2 G), while linolenic acid (18:3 n-3) was the most frequent FA in linseed oil (Figure S2 H). ω-6 FAs (59 %) were predominant in sunflower oil, while linseed oil showed a high ω-3 FA (56 %) content (Table S9) (45).

The lipid droplet phantom experiment served as a control experiment and indicated that FA characteristics ndb (r = 0.762, p < 0.001) and nmidb (r = 0.980, p<0.001) can be adequately quantified independent from the median lipid droplet size (Figure 7B). The carbon chain length (CL) parameter showed a negative correlation between MRS and GC-MS (r = -0.900, p < 0.001). Furthermore, the lipid droplet phantom experiment revealed that the T2 relaxation of the triglyceride’s methylene frequency was highly correlated with the median lipid droplet size (r = 0.988, p < 0.001) independently from the presence of double bounds (Figure 7C). Also, the statistics for T1 relaxation of the triglyceride’s methylene frequency vs. median lipid droplet size showed a negative correlation (r = -0.845, p < 0.001). In contrast, the water component exhibited a positive correlation for the T1 relaxation (r = 0.711, p < 0.001) and a negative more likely exponential correlation for the T2 relaxation (r = -0.849, p < 0.001) vs. the median lipid droplet size, respectively.

**Figure 7:**
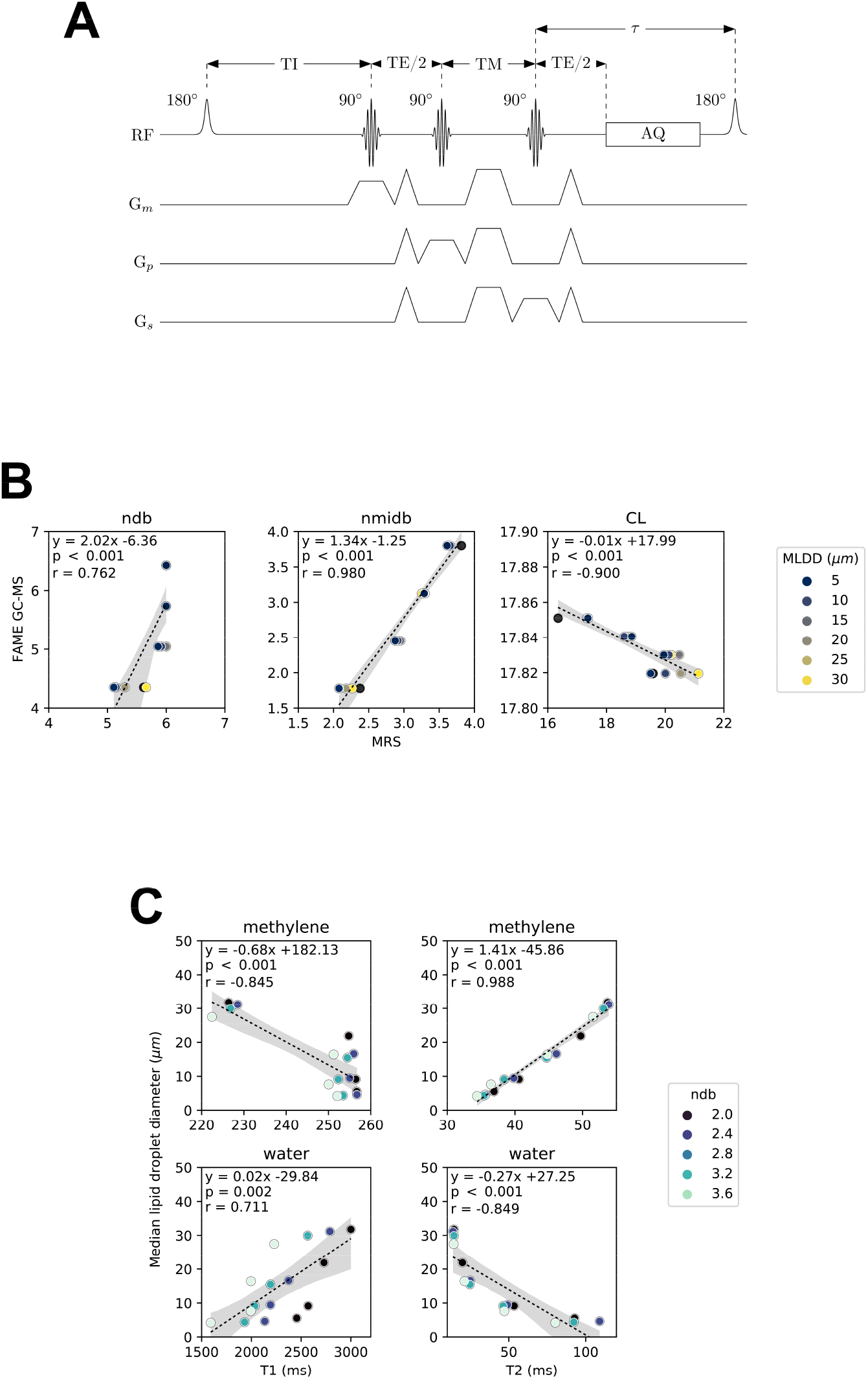
MRS-based characterization of fatty acid composition and lipid droplet size in phantoms. **(A)** MRS pulse sequence diagram of the single-voxel short-TR multi-TI multi-TE (SHORTIE) STEAM sequence. The sequence consists of a regular single-voxel STEAM sequence pattern with a minimal TR (constant recovery delay τ) combined with a non-selective 180-degree inversion RF-pulse. **(B)** Linear regression plots from the phantom experiment: GC-MS-based vs. MRS-based quantification of the FA characteristics ndb, nmidb, CL. The negative correlation for CL (r=-0.900, p<0.001) is considered an artifact arising from the difficulty of it’s accurate modelling and quantification in MRS. MLDD, median lipid droplet diameter. **(C)** Linear regression plots from the phantom experiment of the median lipid droplet diameter vs. the T1 and T2 relaxation of methylene and water, respectively. Very strong correlation between the median lipid droplet diameter and methylene T2 relaxation (r=0.988, p<0.001) independent from the fatty acid unsaturation suggests that methylene T2 is a promising indirect measure of lipid droplet size.

The *in vitro* WAT MRS experiment was conducted with samples from abdominoplasty, general surgery and laparoscopy. Matched MRS, histology and GC-MS data was available for 32 samples originating from 21 donors. Similar to the lipid droplet phantom experiment, a positive but weaker correlation between MRS and GC-MS for the FA characteristics ndb (r = 0.446, p = 0.011) and nmidb (r = 0.773, p < 0.001) was found. No correlation between MRS and GC-MS was observed for FA chain length (Figure 8B).

**Figure 8:**
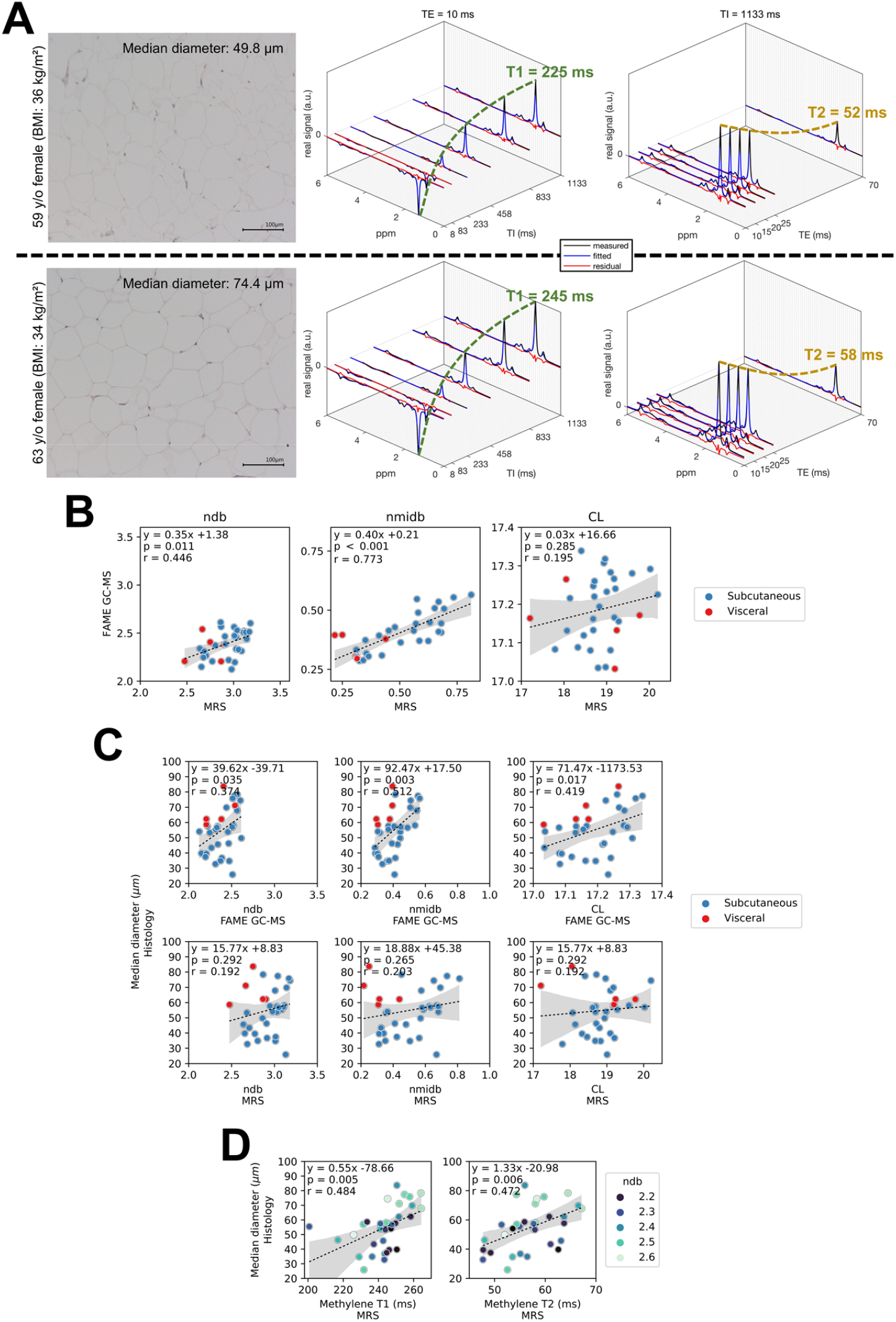
MRS-based characterization of fatty acid composition and fat cell size *in vitro* in adipose tissue samples. **(A)** Exemplary MRS spectra with matched histology for two female subjects with similar age (59 and 63 year) and BMI (36 and 34 kg/m^2^) but difference median fat cell size (49.8 vs.74.4 µm) in abdominal subcutaneous AT. MRS spectra display signal variations with TI (at TE = 10 ms) and TE (TI = 1133 ms), respectively. Fitted methylene T1 and T2 relaxation times are given in green and orange, respectively. **(B)** Linear regression plots from the adipose tissue sample experiment: GC-MS-based vs. MRS-based quantification of the FA characteristics ndb, nmidb, CL. Sample origin is indicated by color. **(C)** Linear regression plots from the adipose tissue sample experiment: GC-MS-based and MRS-based FA characteristics ndb, nmidb, CL vs. median fat cell size. Sample origin is indicated by color. **(D)** Linear regression plot for the methylene T1 and T2 relaxation vs. median fat cell size. Sample ndb (binned) as measured by GC-MS is indicated by color encoding.

The linear regression analysis between the median fat cell size diameter and FA characteristics (Figure 8C) revealed significant moderate correlation for the GC-MS-based ndb (r = 0.374, p = 0.035), nmidb (r = 0.512, p = 0.003) and CL (r = 0.419, p = 0.017), but only showed a trend not reaching significance with the MRS-based ndb (r = 0.192, p = 0.292), nmidb (r = 0.203, p = 0.265) and CL (r = 0.192, p=0.292). In contrast, the median fat cell diameter significantly correlated with both T1 (r = 0.484, p = 0.005) and T2 relaxation (r = 0.472, p = 0.006) of the methylene frequency (Figure 8D).

The histology matched MRS example (Figure 8A) of SAT in comparison from two female donors with comparable age (59 and 63 years) and BMI (36 and 34 kg/m^2^) revealed between the subjects a larger median fat cell size diameter (49.8 vs. 74.4 µm) together with a longer T1 (225 vs. 244 ms) and T2 relaxation time (51 vs. 57 ms) of the methylene frequency.

## DISCUSSION

The depot-specific hypertrophic expansion of adipocytes has been identified as detrimental for the manifestation of metabolic disorders. Understanding the transcriptomic changes occurring with adipocyte hypertrophy and between different WAT depots is therefore of great relevance to identify the causal mechanisms underlying this relationship. Further, the translation of experimental findings into clinical practice remains a challenging task since morphology and composition analysis of WAT are laborious and heavily dependent on biopsy-based techniques. To overcome this limitation, the potential of MRS as a biopsy-free method for the in-parallel assessment of adipocyte size and FA composition was explored.

### SAT and VAT display distinct transcriptomic signatures

Based on the differential expression of multiple developmental genes and transcription factors (*HOX* family, *IRX* family, *BARX1*) our study expands the currently existing body of evidence derived from lineage tracing studies and smaller transcriptional profiling studies indicating that SAT and VAT white adipocytes arise from different progenitors (46–51). Both KEGG gene set enrichment analysis and differential expression of individual genes (e.g. *ALOX15*) highlight that extensive hypertrophic VAT accumulation could lay the transcriptional foundation for the manifestation of metabolic disorders (4, 52–55). As a related finding we observed a significant reduction in the amounts of ARA in VAT. Decreased ARA levels in VAT compared to SAT could stem from an elevated expression of *ALOX15* and other transcripts of enzymes utilizing ARA for the production of pro-inflammatory mediators resulting in an exhaustion of the substrate (40, 56, 57)

### Adipocyte hypertrophy leads to transcriptomic alterations in both SAT and VAT

Amongst other genes, we observed the strongest associations between gene expression and adipocyte size for *EGFL6* and *SLC27A2* in SAT. *EGFL6* has been previously associated with obesity and stimulates the proliferation of adipose tissue-derived stromal vascular cells (43). Further, EGFL6 was found to be differentially expressed with adipocyte size in a recent spatial transcriptomics-based characterization of SAT (13). *SLC27A2* is a FA transporter and an acyl-CoA synthase that shows substrate specificity towards long chain and very long chain ω-3 FAs (58). Expression of *SLC27A2* has previously been shown to be reduced after high fat overfeeding and correlated negatively with obesity and diabetes-related traits (59–61). In VAT a strong negative relationship between genes involved in DNL and elongation of its end products was observed (*FASN*, *ELOVL6*). In good agreement, a strong negative association between saturated FAs with ≤ 20 carbon atoms, typically representing products of DNL with adipocyte size was seen. Decreased rates of DNL could display a physiological response to limit adipocyte expansion and compensate for the increased availability of FAs derived from dietary sources.

Besides alterations in FA metabolism and DNL related genes, negative enrichment for KEGG ribosomal and mitochondrial pathways was observed in relation to adipocyte area in both depots. In contrast, immune and inflammatory pathways i.e. NF-κB signaling were positively enriched and the expression of pro-inflammatory marker genes (*TNF-α* & *IL-6*) was increased in individuals with large adipocytes. Mitochondrial dysfunction due to a reduced expression of respiratory chain genes could i.e. lead to elevated levels of ROS production impacting the endocrine and metabolic functions of the tissue (62). While an elevated production of pro-inflammatory mediators was detected, gene set enrichment analysis suggested negative enrichment for genes involved in ribosomal processes and amino acid synthesis at least partially resembling a senescence associated phenotype (63). Together the identified adipocyte-size associated gene expression patterns provide a transcriptomic foundation for the manifestation of obesity and its related metabolic disorders.

To rule out possible confounding effects of the stromal vascular fraction and different genetic backgrounds mature adipocytes were separated based on size and their transcriptome was characterized using RNA-Seq. Similar pathways and genes were affected in size-separated adipocytes with the large adipocyte fraction displaying a metabolically more harmful expression profile. It is therefore concluded that the detrimental transcriptomic profile of WAT from individuals with large fat cells is at least partially caused by enlarged mature adipocytes themselves. While inflammatory, ribosomal and metabolic pathways were affected in size-separated adipocytes in a similar manner to the cross-sectional analysis on WAT, OXPHOS and thermogenesis did not show significant enrichment. It is therefore hypothesized that OXPHOS gene expression is altered across the entire adipose tissue cell population in individuals with enlarged fat cells. These findings are in excellent agreement with respirometry studies from our group suggesting that mitochondrial respiratory chain capacity is inversely related to BMI and adipocyte size, while no differences in respiratory chain function were observed between size-separated adipocytes (19, 64–66). On the individual gene level, findings on size separated adipocytes are in good agreement with spatial transcriptomics results suggesting that mature adipocytes from different size quartiles display distinct transcriptomic signatures (13).

Significantly less genes were differentially expressed in size-separated adipocytes compared to WAT. While different size fractions from four individuals were compared, the bulk WAT analysis was conducted in more than 140 individuals. Further, in WAT next to mature adipocytes other cell types can contribute to gene expression (67). Another layer of complexity is added as gene expression of adipocytes is dependent on progenitor lineage, spatial localization within the tissue and proximity to other cell types (13).

### Individuals with large adipocytes show a decrease in thermogenic adipocyte content

BAT can dissipate stored chemical energy (triglycerides) in the form of heat mediated by the inner mitochondrial membrane protein *UCP-1*. SAT and VAT browning could display a possible protective mechanism against excessive lipid storage, adipocyte hypertrophy, and at long last metabolic complications. Pathway analysis revealed negative enrichment for lipolysis and FA degradation-related genes in both depots, indicating that hypertrophic adipocytes favor lipid storage over lipid catabolism and thus provide a plausible mechanism for an increase in volume. In parallel, this phenomenon was accompanied by decreased expression of oxidative phosphorylation-related genes, indicating substantial mitochondrial aberrations with adipocyte hypertrophy. As a novel finding, evidence for an inverse relationship between brown/beige adipocyte content and adipocyte size, especially in VAT is provided. A higher thermogenic adipocyte content in individuals with small adipocytes could be protective against excessive lipid accumulation, adipocyte hypertrophy and its associated metabolic complications. Vice versa, brown and beige adipocyte content seems to be blunted in individuals with large adipocytes giving a possible explanation for disproportionate lipid storage, an increased risk for adipocyte hypertrophy and an adverse transcriptomic and metabolic profile. As our study is associative, whether genetic predisposition, environmental stimuli or a bidirectional effect lead to the observed inverse relationship remains elusive and should be the object of future longitudinal studies. Together, with reduced expression of lipolytic and FA degrading genes our data indicate that a transcriptional shift from an energy-burning towards a lipid storing, pro-inflammatory phenotype is a central hallmark of hypertrophic WAT.

### MRS-based methods for the simultaneous characterization of WAT morphology and FA composition

Being able to determine WAT FA composition and adipocyte size in parallel as prognostic biomarkers for metabolic disease using non-invasive diagnostic tools would be highly desirable. Therefore, an MR-based technique for the simultaneous characterization of fat cell size and FA composition was developed to pave the way towards a non-invasive assessment of WAT morphology and composition that could be applicable in a clinical context.

The lipid droplet phantom experiment demonstrated the feasibility of the characterization of the FA characteristics as described by ndb and nmidb, as previously reported (38). Beyond what was already known, the experiment empirically demonstrated for the first time the correlation of the methylene T2 relaxation with median lipid droplet size independent from the presence of double bounds in the considered range. Thus, the methylene T2 relaxation can be used as an indirect measure of lipid droplet size. As mature white adipocytes are unilocular with the lipid droplet taking up most of the intracellular space lipid droplet size characterization by T2 relaxation times can be considered representative for the estimation WAT fat cell size. With MR relaxation properties being temperature-dependent it should be acknowledged that *in vivo* relaxation times at body temperature will differ from the reported relaxation times of excised room temperature WAT samples. Interestingly, SHORTIE MRS-based ndb and nmidb measures correlated well with those from GC-MS, but not with the median adipocyte diameter whereas SHORTIE MRS-based ndb and nmidb correlated with the GC-MS-based derivatives of ndb and nmidb. While we found a similar trend of increasing methylene T2 relaxation times with increasing lipid diameter in WAT, the signifcance of the correlation flipped for methylene T1 relaxation showing an increase in T1 with increasing diameter. The differences between the phantom and the WAT sample experiment maybe partly explained in consideration of a) the performed GC-MS and MRS are not characterizing the exact same pool of fatty acids and b) structural difference between emulsified lipid droplets and WAT microstructure. The relaxation-based characterization of the median fat cell size in WAT has several advantages compared to approaches based on the probing of restricted diffusion effects including reduced data acquisition times, reduced sensitivity to vibrational artefacts and reduced motion sensitivity (17, 18, 68). Furthermore, the reported MR parameter relationship pinpoints the possibility of a methodological translation of the proposed measurement framework into a quantitative imaging-based method allowing the spatially-resolved probing of multiple AT depots.

Therefore, an application towards an unperturbed in-vivo phenotyping of adipocyte size and FA composition is feasible and the object of future investigations. Being able to simultaneously probe morphological and compositional parameters of WAT in a non-invasive manner would offer the opportunity to longitudinally monitor adipocyte size and FA composition in various depots of the body. Further, parallel MR-based measurements of FA patterns and adipocyte hypertrophy could serve as a potential biomarker to detect early changes in WAT morphology and composition that might occur prior to the systemic manifestation of metabolic diseases.

Despite validating our findings from bulk tissue in size-separated adipocytes we cannot fully rule out a substantial contribution of other cell types from the stromal vascular fraction to the observed changes in gene expression in bulk tissue characterized by adipocyte hypertrophy. The reported MRS relaxation times could be also affected by other tissue properties beyond adipocyte size (e.g. tissue oxygenation) (69). The employed MRS sequence was not derived from a formal optimal design of experiment and could therefore be further optimized. J-modulations were considered neglectable and not accounted for in the signal modeling process which, however, may further improve FA characterization irrespective of its computational demand and modelling uncertainties.

The observed switch to an energy storing, pro-inflammatory transcriptome in individuals with large adipocytes, especially in metabolically more harmful VAT provides important novel mechanistic insights on the transcriptional background of hypertrophic obesity. The identified reduction in thermogenic adipocyte content in individuals with large fat cells could display a novel mechanism facilitating hypertrophic WAT expansion and manifestation of metabolic disorders. Our MR-based proof-of-concept approach could become a promising tool for the non-invasive estimation of FA composition and adipocyte size in-vivo. An earlier and better diagnosis of metabolic derangements based on adipose tissue morphology and FA composition before systemic manifestations occur could substantially improve the treatment and prevention of cardiometabolic disorders. The present work is a first step towards this goal, but needs confirmation in prospective human studies.

## ACKNOWLEDGEMENTS

We are grateful to Prof. Dr. T. Huettl, Dr. U.Schulze-Eilfing, Dr. C. Kleeberger, Dres. Hoffmann, Dr. O. Prokopchuk and their respective teams for obtaining adipose tissue samples. We thank S. Winkler for assisting with WAT sample collection, D. Weidlich for helpful discussions, M. Zamskiy, C. Held and L. Patzelt for helping with the logistics and execution of the MRS measurements.

## SUPPLEMENTARY INFORMATION

**Figure S1:**
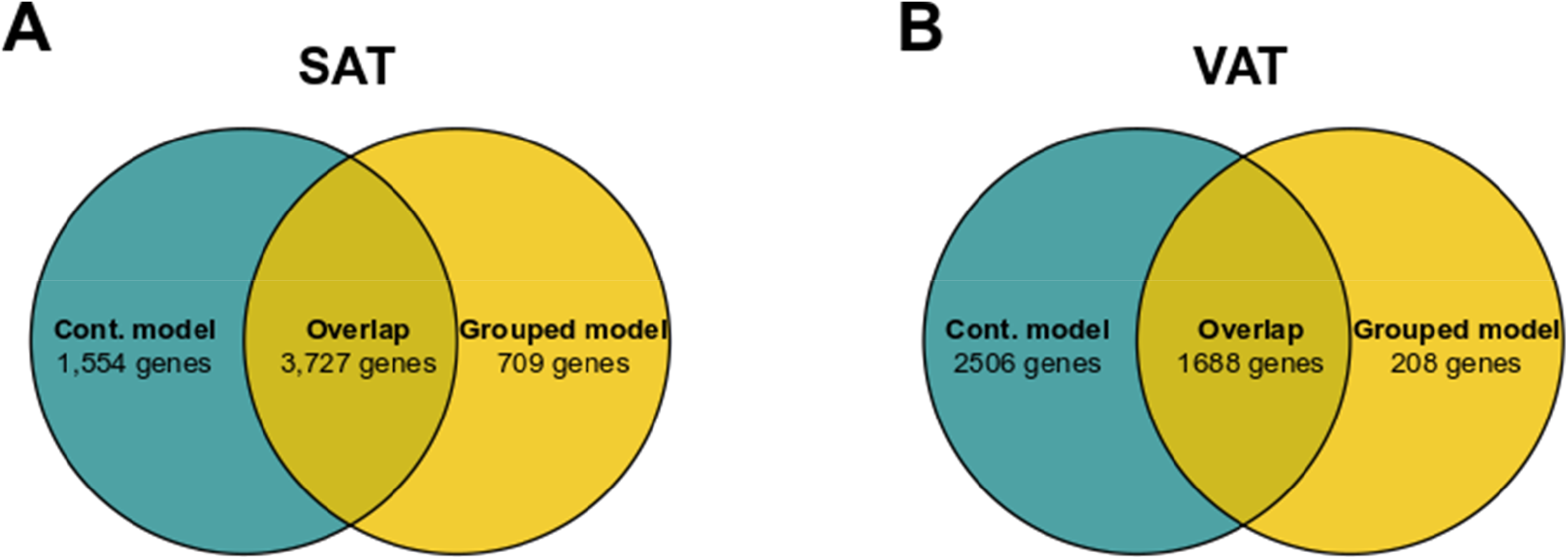
Number of differentially expressed genes in relationship to the applied differential expression model in GTEx samples (continuous vs. grouped) (A) In SAT 1,554 genes were unique for the continuous model (turquoise) while 709 genes were differentially expressed solely in the grouped model (yellow). Both models shared significant differential expression (FDR < 0.05) of 3,727 genes. (B) In VAT 2,506 genes were unique for the continuous model (turquoise) while 208 genes were differentially expressed solely in the grouped model (yellow). Both models shared significant differential expression (FDR < 0.05) of 1,688 genes.

**Figure S2:**
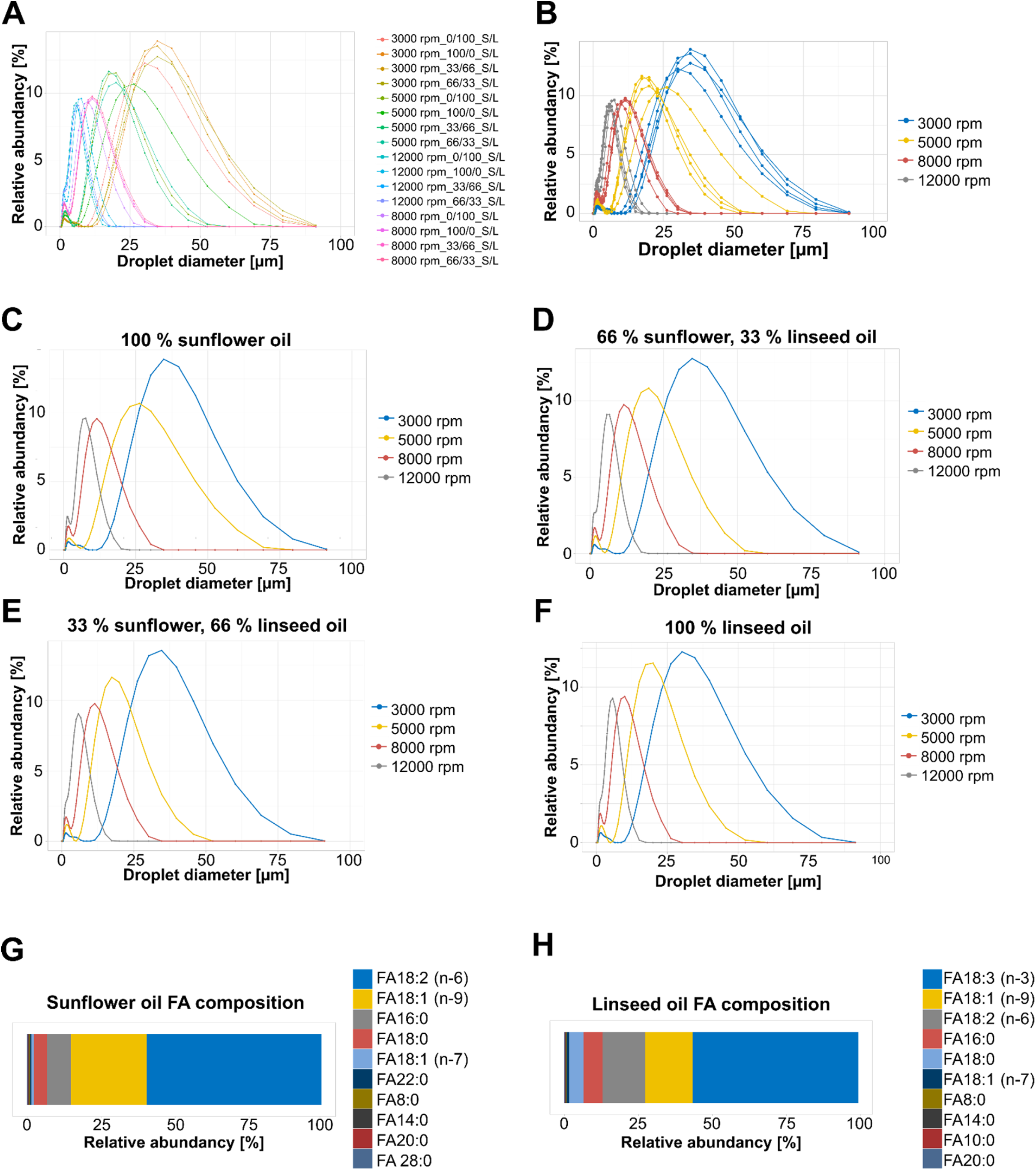
Characterization of water-fat phantoms (A) Density plots from all water-fat phantoms. (B) Density plots from all water-fat phantoms colored according to stirrer rpm. (C) Density plots from the 100 % sunflower oil water-fat phantoms. (D) Density plots from the 66 % sunflower 33 % linseed oil water-fat phantoms. (E) Density plots from the 33 % sunflower 66 % linseed oil water-fat phantoms. (F) Density plots from the 100 % linseed oil water-fat phantoms. (G) Top ten most abundant fatty acid species in sunflower oil. (H) Top ten most abundant fatty acid species in linseed oil.

**Table S5:** Adipocyte diameter and area of FAME GC-MS samples

|  | SAT<br>FAME GC-MS | VAT<br>FAME GC-MS |
| --- | --- | --- |
| n | 22 | 12 |
| Mean Diameter<br>±<br>SD [µm] | 3,457<br>±<br>1638 | 3,715<br>±<br>1070 |
| Diameter range<br>min – max [µm] | 1,307 – 6,410 | 2,092 – 5,544 |
| Mean Area<br>±<br>SD [µm] | 59.59<br>±<br>15.36 | 63.63<br>±<br>9.23 |
| Area range<br>min – max [µm] | 36.74 – 85.92 | 48.23 – 77.06 |

**Table S6:**
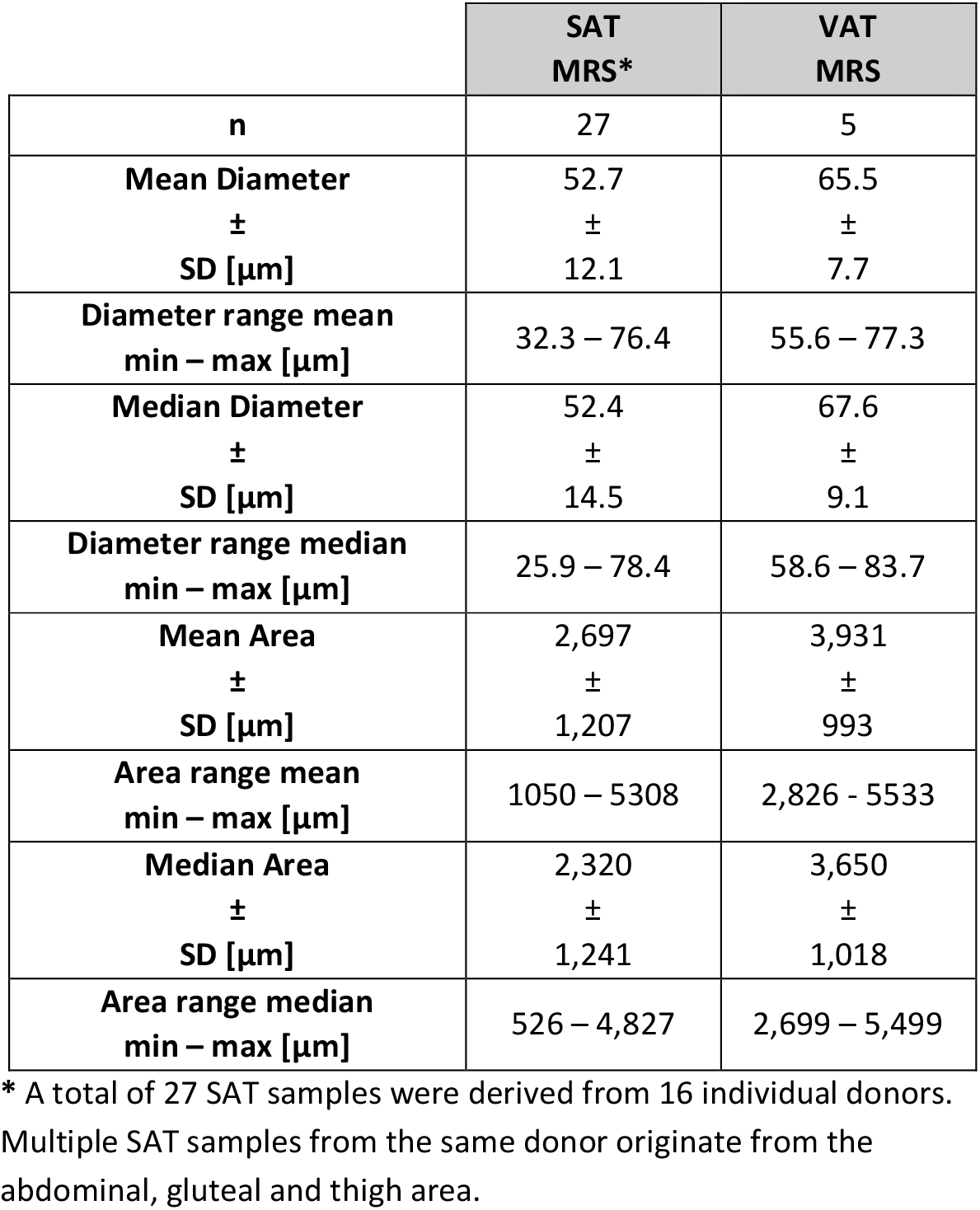
Adipocyte diameter and area of MRS samples

|  | SAT<br>MRS* | VAT<br>MRS |
| --- | --- | --- |
| n | 27 | 5 |
| Mean Diameter<br>±<br>SD [µm] | 52.7<br>±<br>12.1 | 65.5<br>±<br>7.7 |
| Diameter range mean<br>min – max [µm] | 32.3 – 76.4 | 55.6 – 77.3 |
| Median Diameter<br>±<br>SD [µm] | 52.4<br>±<br>14.5 | 67.6<br>±<br>9.1 |
| Diameter range median<br>min – max [µm] | 25.9 – 78.4 | 58.6 – 83.7 |
| Mean Area<br>±<br>SD [µm] | 2,697<br>±<br>1,207 | 3,931<br>±<br>993 |
| Area range mean<br>min – max [µm] | 1050 – 5308 | 2,826 - 5533 |
| Median Area<br>±<br>SD [µm] | 2,320<br>±<br>1,241 | 3,650<br>±<br>1,018 |
| Area range median<br>min – max [µm] | 526 – 4,827 | 2,699 – 5,499 |

\* A total of 27 SAT samples were derived from 16 individual donors. Multiple SAT samples from the same donor originate from the abdominal, gluteal and thigh area.

**Table S7:**
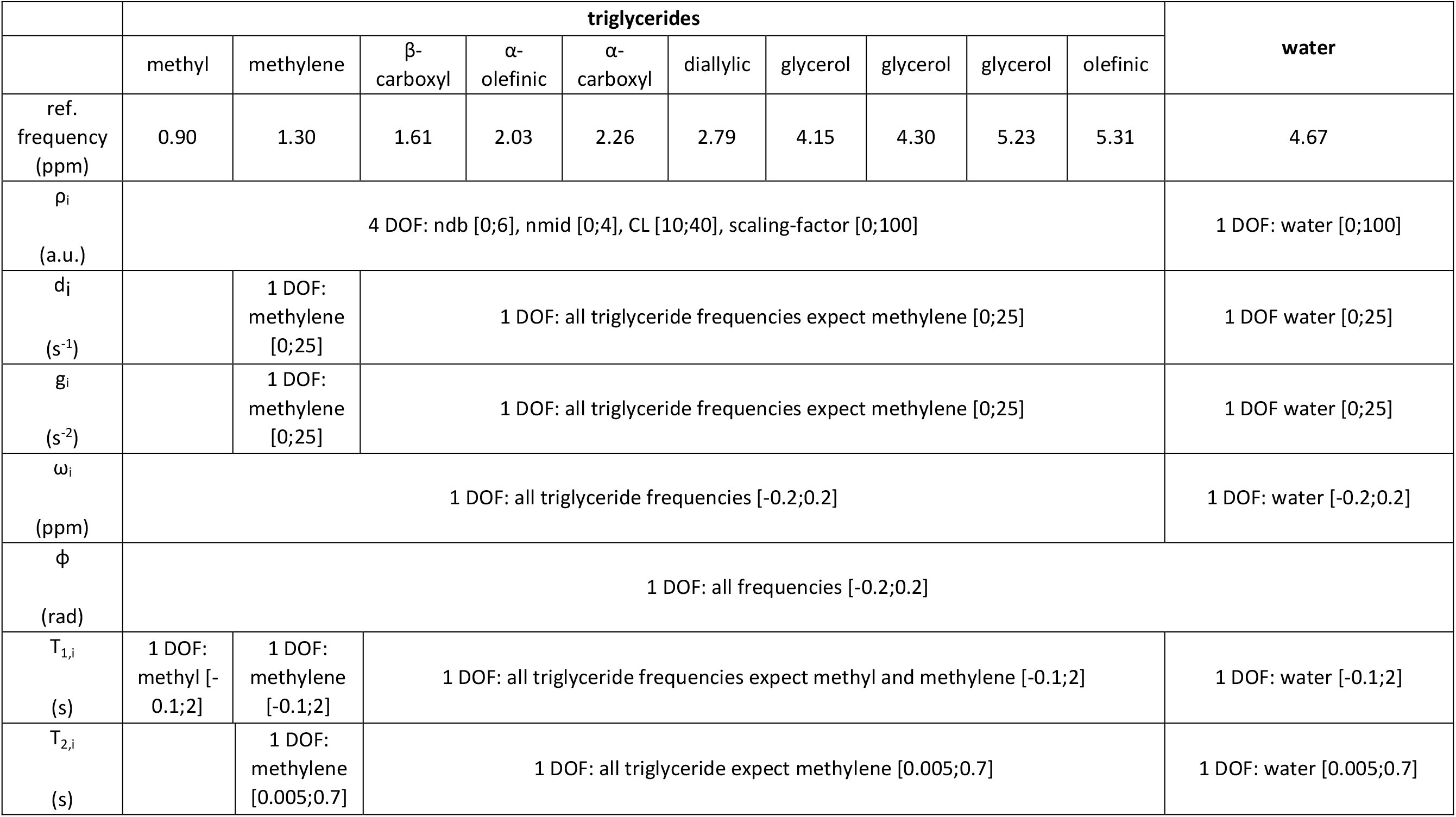
MRS signal fitting model consisting of in total eleven frequency components: 10 triglyceride frequencies and 1 water frequency. Using triglyceride-model-constraints and relaxation constraints for the fat frequencies lead to a reduction of model parameters resulting in a total of 21 degrees of freedom (DOF). Parameter bounds are given in square brackets, e.g. [lower bound; upper bound]. a.u., arbitrary unit.

**Table S8:** Lipid droplet sizes of water-fat phantoms

| Revolutions per minute<br>[rpm] | Mixing ratio<br>Sunflower/Linseed | Diameter<br>First decile<br>[μm] | Diameter<br>Median<br>[μm] | Diameter<br>Ninth decile<br>[μm] |
| --- | --- | --- | --- | --- |
| 3000 | 100/0 | 17.31 | 31.80 | 50.76 |
|  | 66/33 | 16.34 | 31.19 | 51.94 |
|  | 33/66 | 16.42 | 29.97 | 48.50 |
|  | 0/100 | 13.93 | 27.48 | 46.64 |
| 5000 | 100/0 | 8.28 | 21.88 | 39.54 |
|  | 66/33 | 3.37 | 16.62 | 29.86 |
|  | 33/66 | 3.17 | 15.45 | 26.85 |
|  | 0/100 | 5.56 | 16.41 | 28.59 |
| 8000 | 100/0 | 1.65 | 9.14 | 17.47 |
|  | 66/33 | 1.68 | 9.46 | 17.92 |
|  | 33/66 | 1.62 | 9.10 | 17.20 |
|  | 0/100 | 1.40 | 7.62 | 14.71 |
| 12000 | 100/0 | 1.19 | 5.54 | 10.47 |
|  | 66/33 | 1.19 | 4.63 | 9.03 |
|  | 33/66 | 1.21 | 4.36 | 8.57 |
|  | 0/100 | 1.19 | 4.20 | 8.12 |

**Table S9:** Fatty acid composition of sunflower and linseed oil

| Sample | ndb | nmidb | CL | SFA<br>[0-1] | UFA<br>[0-1] | MUFA<br>[0-1] | PUFA<br>[0-1] | n3<br>[0-1] | n6<br>[0-1] |
| --- | --- | --- | --- | --- | --- | --- | --- | --- | --- |
| Sunflower | 4.35 | 1.78 | 17.82 | 0.14 | 0.86 | 0.27 | 0.59 | 6.99e<sup>-04</sup> | 0.59 |
| Linseed | 6.43 | 3.80 | 17.85 | 0.12 | 0.88 | 0.17 | 0.70 | 0.56 | 0.14 |

## Notes

### Competing Interest Statement

The authors have no conflicts of interest to disclose. C.M.L. has collaborated with Novo Nordisk and Bayer in research, and under a university agreement, did not accept any personal payment. M.C. has collaborated with Bayer without accepting personal payment. M.C. further serves as a member on the Scientific Advisory Board of Nestle and holds equity in the company Waypoint Bio.

